# Phospholipase C6 Regulates Hydroxyproline *O*-arabinosyltransferase-mediated Pollen Tube Growth in *Arabidopsis*

**DOI:** 10.1101/2022.10.26.513895

**Authors:** Steven Beuder, Xuesi Hua, Alexandria Dorchak, Cora A. MacAlister

## Abstract

Hydroxyproline *O*-arabinosylation is a highly-conserved and plant-specific post-translational modification found on extensins and other structural proteins in the cell wall, and is catalyzed by Hydroxyproline *O-*arabinosyltransferases (HPATs). In *Arabidopsis*, loss of *HPAT1* and *HPAT3 (hpat1/3*) causes reorganization of components in the pollen tube (PT) cell wall, which compromises cell wall structural integrity and decreases PT growth and fertility. We have previously shown that reduced secretion (caused by loss-of-function mutations in secretory genes *EXO70A2, SEC15A*, and *SEC1A*) suppressed cell wall defects and strongly rescued poor growth and fertility in *hpat1/3* PTs. Here, we show that a missense mutation in *PHOSPHOLIPASE C6 (PLC6*) also rescues *hpat1/3* PT growth and fertility. Transgenic insertion mutations that disrupt *PLC6* expression did not improve *hpat1/3* pollen fertility, and did not affect PT growth or fertility in the wild type background. This data suggests that our missense allele (*plc6-4*) does not function like a true loss-of-function allele, and that PLC6 is not required for wild type PT growth. However, in the absence of *hpat1/3, plc6-4* PTs have defects in transmission and polarized growth, as indicated by meandering growth paths and a resulting crooked appearance. *plc6-4* PT elongation and straightness are more sensitive to elevated levels of calcium than wild type. This may be due the nature of the *plc6-4* mutation, which causes an E569K amino acid substitution in the lipid-binding C2 domain. The 569 position is located among conserved residues that bind calcium. The resulting charge inversion caused by the E569K substitution may disrupt PLC6’s lipid binding and phospholipase activities. Here, we show that *PLC6* influences polarized PT growth and HPAT-mediated PT growth and fertility, and future studies are necessary to better understand the relationship between calcium and *PLC6* in PT growth.

## 1. Introduction

Pollen tubes (PTs) elongate through a highly polarized growth mechanism called tip growth, which relies on the careful regulation of structural and mechanical properties of the cell wall. For proper tip growth, PTs must maintain an extensible apical tip to allow for elongation, and rigid subapical walls to prevent multidirectional expansion (leading to PT swelling or branching) and premature bursting. In order to maintain two spatially distinct regions of the cell wall with different mechanical properties, PTs differentially regulate the structures of these two wall regions; however, the molecular mechanisms underlying this process are still unclear.

The plant cell wall is a complex extracellular matrix composed of multiple carbohydrate networks including cellulose, hemicellulose, callose and pectins, as well as proteins such as cell wall-remodeling enzymes and glycoproteins. Pectins have a major influence on the mechanical properties of the cell wall, and mutations in genes that control pectic structure severely impair tip growth and pollen fertility, leading to decreased seed production and fitness (Jiang et al., 2005; Leroux et al., 2015; Röckel et al., 2008). Pectins are a family of structurally diverse carbohydrates, with homogalacturonan (HG) being the most abundant polymer type (Mohnen, 2008). HG is composed of *alpha-*1,4-linked-d-galacturonic acid (GalA) and is highly methylesterified (me-HG) by enzymes in the Golgi prior to secretion (Krupková et al., 2007; Mohnen, 2008; Mouille et al., 2007; O’Neill et al., 1990; Ridley et al., 2001). After secretion into the cell wall, pectin methylesterases catalyze the de-methylesterification of me-HG to form dme-HG, exposing negatively charged carboxyl groups. Neighboring dme-HG molecules then undergo crosslinking interactions via calcium bridges (described as the “egg-box” model), which promotes gelation and increases the overall rigidity of the cell wall (Grant et al., 1973; Pelloux et al., 2007). In PTs, me-HG is localized to the cell wall and enriched at the tip, while dme-HG localizes to the subapical walls and is absent from the tip (Chebli et al., 2012).

Pectins also interact with extensins, which are a family of hydroxyproline (Hyp)-rich glycoproteins that regulate the formation and structural integrity of the cell wall (Choudhary et al., 2015; Hall & Cannon, 2002). The structures of “classical” extensins are amphiphilic and highly repetitive, consisting primarily of alternating Ser(Pro)3-5 motifs and Tyr/Tyr-Val motifs, and are also typically rich in Lys, giving them an overall positive charge (Showalter et al., 2010). Tyr containing motifs facilitate intermolecular crosslinking of extensin monomers to form networks, and negatively charged pectic groups interact with positively charged extensins to form extensin pectate (Cannon et al., 2008; Smith et al., 1984). Through these interactions, it has been proposed that extensin networks function as a scaffold for pectin assembly and possibly regulate crosslinking of dme-HG to influence cell wall mechanics (Cannon et al., 2008; MacDougall et al., 2001).

Extensins undergo significant post-translational modifications prior to secretion into the cell wall. Prolines found in Ser(Pro)3-5 motifs are hydroxylated by prolyl 4-hydroxylases (P4Hs) to form Hyp (Tiainen et al., 2005), which are then *O-*arabinosylated by a cascade of Golgi-localized glycosyltransferases to form linear oligoarabinosides that are typically 3-5 residues long (Akiyama et al., 1980; Petersen et al., 2021). The first arabinose is added by enzymes called Hyp *O-*arabinosyltrasferases (HPATs), which are encoded by a three member gene family in *Arabidopsis* (Ogawa-Ohnishi et al., 2013). Loss of *HPAT1* and *HPAT3 (hpat1/3*) drastically decreases seed production due to poor growth and decreased fertility of *hpat1/3* PTs (MacAlister et al., 2016). In addition to poor elongation, *hpat1/3* PTs burst more frequently and exhibited morphological defects including increased widths and the initiation of secondary, sub-apical tips (branching) compared to wild type (WT), consistent with a loss of structural integrity of the cell wall. Through immunolabeling and other staining techniques, we identified and characterized multiple differences in the cell wall structure of *hpat1/3* PTs compared to WT (Beuder et al., 2020). One notable difference was that more dme-HG signal was detected throughout the cell wall of *hpat1/3* PTs, including at the tip where it is usually absent (Chebli et al., 2012), suggesting that overall rigidity is increased in *hpat1/3* PT cell walls and tip extensibility is decreased (Beuder et al., 2020).

To learn more about how HPATs regulate PT growth, we wanted to identify additional genes involved in this pathway. We performed a mutagenesis screen and identified plants in which the *hpat1/3* PT defects were suppressed, and mapped the *hpat1/3* suppressor mutations using whole-genome sequencing (described in Beuder & MacAlister, 2020). We previously characterized one *hpat1/3* suppressor mutant called *fertilization restored in hpat1/3 1* (*frh1*), in which the suppression-causing mutation was identified as a G319E missense mutation in *EXO70A2 (exo70a2-2*), which encodes an EXO70 isoform (Beuder et al., 2020). EXO70 is a member of the exoycst complex, which facilitates tethering of secretory vesicles to the plasma membrane prior to SNARE-mediated membrane fusion and exocytosis. *hpat1/3* PT growth defects were strongly rescued in the *hpat1/3; exo70a2-2/frh1* suppressor, and dme-HG immunolabeling and other structural defects were also partially suppressed. Furthermore, apical secretion of a GFP-based reporter with an EXTENSIN 3 (EXT3) Ser(Pro)3-5 motif [GF(EXT3)P] was increased in *hpat1/3* PTs. This was rescued/decreased in *exo70a2* mutants, suggesting that EXO70A2 promotes secretion of extensins and extensin-like proteins. The GF(EXT3)P reporter is Hyp *O-*arabinosylated in an HPAT-dependent manner; therefore, our data suggested that Hyp *O-*arabinosylation may regulate the rate of EXO70A2-mediated extensin secretion. We also observed that loss of function mutations in *SEC15A* suppressed *hpat1/3* PT growth defects (Beuder et al., 2022). Sec15 is another member of the exocyst complex and is required for PT germination, growth, and fertility (Hála et al., 2008). Taken together, our findings indicated that that global knockdown of exocyst-mediated secretion complements the cell wall defects caused by *hpat1/3* (Beuder et al., 2020).

*hpat1/3* pollen fertility defects were also strongly suppressed by multiple independent mutations in *SEC1A*, which encodes a Sec1/Munc18 protein that regulates the formation of SNARE complexes between vesicle and target membranes prior to fusion (Beuder et al., 2022; Südhof & Rothman, 2009). Sec1a and Keule (a Sec1p ortholog) function redundantly in pollen and are required for PT germination and fertility (Beuder et al., 2022). *sec1a-1* strongly rescued PT growth and seed set in *hpat1/3*, and also decreased GF(EXP3)P secretion and pollen germination, similar to the effects caused by *exo70a2*. In summary, these findings demonstrated that structural properties of the cell wall are regulated by a novel relationship between the secretory pathway and Hyp *O*-arabinosylation of extensins and extensin-like proteins, which must be carefully balanced to promote proper PT growth and fertility. Here, we describe another *hpat1/3* plant family (*frh3*) with improved pollen fertility caused by a mutation mapped to *PHOSPHOINOSITIDE (PI)-SPECIFIC PHOSPHOLIPASE C6 (PLC6). PLC6* (At2g40116) is one of nine PLCs (*PLC1* through *9*) encoded in the genome of *Arabidopsis thaliana*.

PLCs have known roles in secretion and PT growth by regulating phosphoinositide availability through phospholipase activity. PLCs bind and cleave phosphatidylinositol 4,5-bisphosphate [PI(4,5)P_2_] to form the secondary messengers diacylglycerol (DAG) and inositol trisphosphate (IP3). EXO70 binds PI(4,5)P_2_ at the plasma membrane to promote exocyst binding and vesicular fusion (He et al., 2007; Pleskot et al., 2015). PI(4,5)P_2_ was observed to localize to the plasma membrane at the tip of tobacco PTs and *Nicotiana tabacum* (Nt) PLC3 localized to the subapical walls, suggesting that NtPLC3-mediated PI(4,5)P_2_ hydrolysis at these regions confines PI(4,5)P_2_ to the tip (Helling et al., 2006). PI(4,5)P_2_s are formed by the phosphorylation of PIP4 by PIP4 5-kinases. Knockdown of PI4P 5-kinase activity decreases PT growth rates and germination frequency, and PI4P 5-kinase overexpression increases pectin accumulation and thickens the cell wall of PT tips tip (Ischebeck et al., 2008). In addition to mediating secretion, PI(4,5)P_2_ signaling is important for other cellular processes. In plants, IP3 is quickly converted to IP6 which triggers the release of intracellular calcium stores to increase cytoplasmic calcium concentration (Lemtiri-Chlieh et al., 2003). PTs maintain a tip-focused calcium gradient which controls growth direction (Herbell et al., 2018; Taiz, 1984), but it is unclear if, or how, this is mediated though PLC-PI(4,5)P_2_ signaling.

Much of what we know about PLC-PI signaling comes from animal systems, and plants lack some of animals’ signaling machinery. For example, plant PLCs lack an N-terminal pleckstrin homology (PH) domain, which directly binds PI(4,5)P_2_. PI(4,5)P_2_ levels are also particularly low in plant cell plasma membranes, suggesting that there may be other endogenous PLC ligands *in planta*. Plant PLC structures include a variable EF hand domain at the N-terminus, followed by conserved X and Y catalytic domains and a C-terminal C2 lipid binding domain (Munnik & Testerink, 2009). For one rice PLC (AK064924), the C2 domain alone was sufficient to bind lipids in a calcium-dependent manner (Rupwate & Rajasekharan, 2012). However, NtPLC3 required both the EF and C2 domains to bind the plasma membrane in tobacco PTs (Helling et al., 2006).

Calcium is an important regulator of PLC function, as it is required for phospholipase activity and lipid binding (Munnik & Testerink, 2009; Rupwate & Rajasekharan, 2012), and the C2 domain harbors several important calcium biding regions (Rebecchi & Pentyala, 2000). When the C2 domain binds calcium, the protein undergoes a conformational change that stabilizes the active site residues in an orientation for catalysis (Wang, 2001), and exposes hydrophobic surfaces to promote lipid binding (Rupwate & Rajasekharan, 2012).

Here, we interrogate the potential links between *PLC6* and the secretory pathway in HPAT-mediated PT growth, and show that PLC6 regulates PT polarity and growth response to calcium.

## 2. Materials and Methods

### 2.1 Plant growth conditions and suppressor screen

Columbia-0 WT *Arabidopsis* and mutants were grown under 16 hour light/ 8 hour dark conditions in controlled growth room maintained at 23° C. Mutagenesis of *hpat1/3* seeds, suppressor phenotyping and selection, whole-genome sequencing and analysis steps were performed as previously described in (Beuder et al., 2020; Beuder & MacAlister, 2020).

### 2.2 *plc6-4* genotyping

Derived Cleaved Amplified Polymorphic Sequences (dCAPS) primers (Supplemental Table 1) were designed to detect the *plc6-4* SNP in the *frh3* genomic DNA. Primers were designed to create a full-length PCR product of 207 base pairs, and the forward dCAPS primer contained a mismatch that created a recognition site for the restriction enzyme Hyp188III only when the SNP was also present in the PCR product. Therefore, the WT sequence (containing guanine) would not be cut, while the mutant sequence (containing adenine) would be cut to generate two fragments of sizes 186 bp and 21 bp, which were detected through gel electrophoresis with a 2-3% agarose gel.

### 2.3 PT assays and seed counts

All pollen germination and growth assays were carried out using *in vitro* pollen growth media (PGM) (modified from Rodriguez-Enriquez et al., 2013) consisting of 10% sucrose, 0.01% boric acid, 1mM CaCl_2_, 1mM Ca(NO_3_)_2_, 1mM KCl, 0.03% casein enzymatic hydrolysate, 0.01% myo-inositol, 0.1 mM spermidine, 10mM γ-Aminobutyric acid, 500μM methyl jasmonate, pH adjusted to 8.0 and solidified with 1% low melting temperature agarose. For experiments with varying calcium concentrations the molar ratio between CaCl_2_ and Ca(NO_3_)_2_ was maintained while increasing or decreasing to the appropriate level. PTs were grown on cellophane placed on top of the semi solid media in plates, which were incubated in a homemade humid chamber (glass box with wet paper towls) in the dark. Length, width, germination, and bursting frequency assays were performed as previously described (Beuder et al., 2020).

For seed counts, mature siliques (just starting to turn yellow) were plucked and placed in 70% ethanol for several days to remove pigment. Once the seeds were clearly visible through the cleared silique, seeds were counted using a dissecting microscope.

### 2.4 Cloning

The genomic suppressor rescue construct (*PLC6p:PLC6*) was cloned by amplifying the *PLC6* gene sequence (2530 base pairs) and the entire upstream region before the end of the previous gene (875 base pairs) using the Phusion^®^ High-Fidelity DNA Polymerase (New England Biolabs M0530S) from WT Col-0 genomic DNA extracted from leaf tissue. Primers used are listed in Supplemental Table 1. *PLC6:PLC-mNeonGreen* (mNG) was cloned by first amplifying the same region with a different reverse primer to omit the stop codon, and then amplifying the mNG sequence from plasmid DNA. We fused mNG to the *PLC6* C-terminus using the Gateway cloning system. Recombinant sequences were cloned into the binary vector pFAST-G01 (*PLC6p:PLC6*) or pFAST-R01 (*PLC6p:PLC6-mNG*) (Shimada et al., 2010) and transformed into plants with Agrobacterium.

### 2.5 Microscopy

PTs were imaged for growth, bursting, germination, and morphological assays with a Leica DM5500 compound microscope with DIC optics at 10X magnification. GF(EXT3)P+ PTs were imaged using the same microscope but with a GFP filter. For imaging of PLC6-mNG fusions, PTs were grown *in vitro* on slides with rubber spacers containing PGM for 2 hours in a humid chamber in the dark. After growth time, liquid PGM with 4 μM FM4-64 was added on top of the PTs and covered with a coverslip. PTs were imaged using a Leica SP5 laser-scanning confocal microscope 10 minutes after FM4-64 application. To image PLC6-mNG, we used an excitation laser with 488 nm wavelength, a RSP500 dichroic beam splitter, and detectors were set to capture light with a wavelength range of 494-575 nm. To image FM4-64, we used an excitation laser with 514 nm wavelength, a DD 458/514 dichroic beam splitter, and detectors were set to capture light with a wavelength range of 620-783 nm. Images were overlayed with ImageJ.

### 2.6 Aniline blue staining

Aniline blue staining and imaging was performed similarly as described in Lara-Mondragón & MacAlister, 2021, with several changes. WT flowers were emasculated and manually pollinated with WT or *plc6-4* pollen and fixed after 24 hours. Pistils were transferred to an Eppendorf tube and vacuum infiltrated with 1 mL acetic acid:ethanol (1:3) for 2 hours, and then transferred to 5M NaOH overnight. Pistils were transferred to Aniline Blue Fluorochrome (Biosupplies Catalog Number 100-1) (0.001 mg/ml in 0.1 M K_2_HPO_4_, pH 10), mounted in VectaShield (Vector Laboratories, H-1400), and imaged with a Leica DM5500B microscope using a DAPI filter (355/455 nm) and a Leica DFC365FX camera.

## 3. Results

### 3.1 *frh3* suppresses *hpat1/3* PT growth and fertility defects to improve seed production

WT *Arabidopsis* plants typically produce large siliques filled with seeds due to efficient self-fertilization of all or most ovules. Self-fertilized *hpat1/3* plants produce fewer seeds due to poor growth of *hpat1/3* PTs, which drastically decreases fertilization efficiency (MacAlister et al., 2016). Through a mutagenesis screen (described in Beuder et al., 2020; Beuder & MacAlister, 2020), we identified *hpat1/3* plants (*frh3*) with significantly increased seed set, and no observable abnormal vegetative phenotypes (Figure 1A-C).

**Figure 1.**
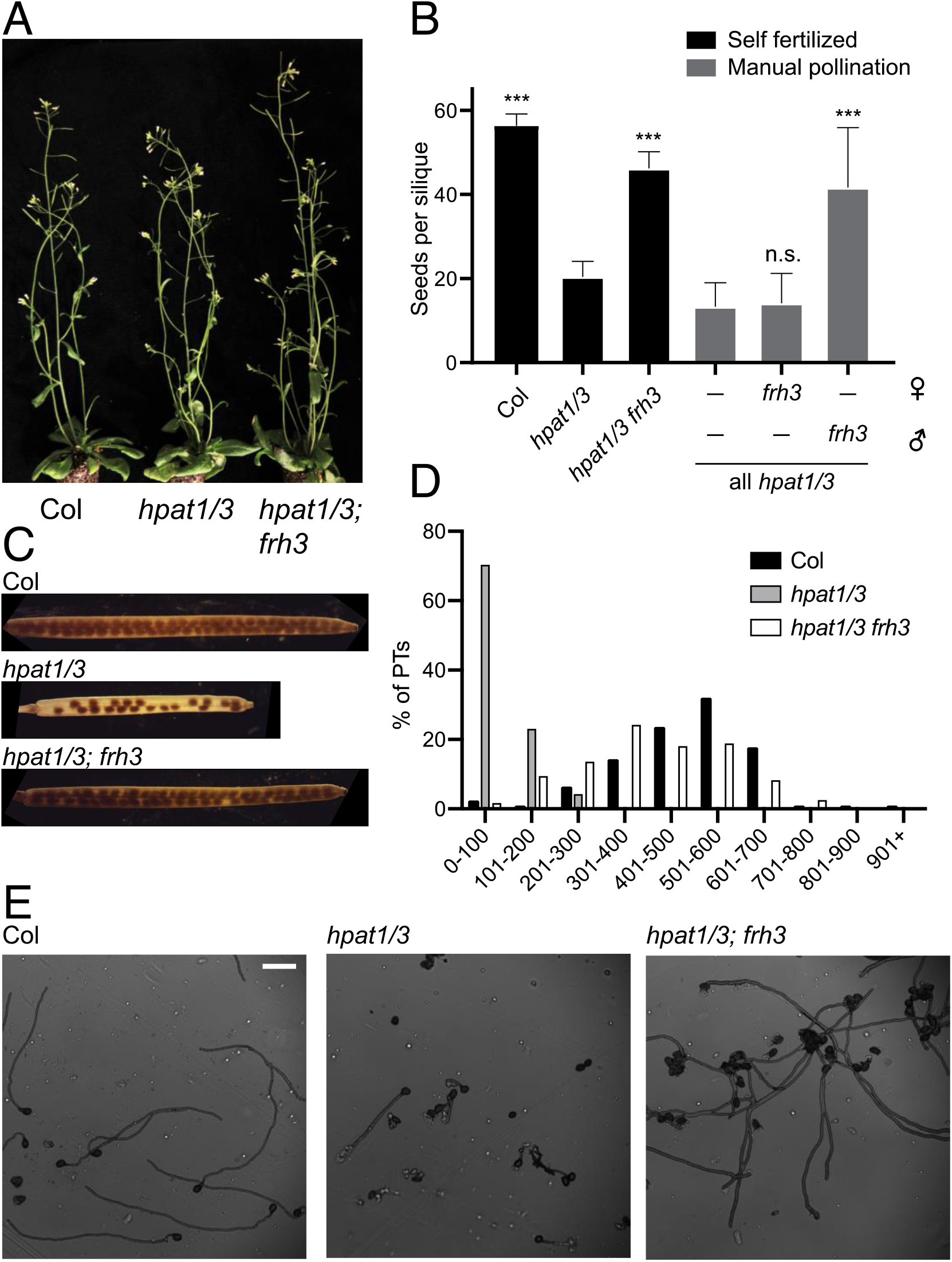
*hpat1/3* pollen fertility defects and seed production are rescued in frh3. A) *hpat1/3* and *hpat1/3; frh3 (frh3*) plants show no vegetative abnormalities. B) Seed counts (number of seeds per silique) for WT, *hpat1/3* and *frh3*. Bars represent average seed counts and error bars represent standard deviation. (N ≥ 10 siliques per genotype). Statistical analysis performed using Student’s t-test. *** denotes p-value ≤ 0.0005. C) Representative siliques from each genotype cleared in 70% ethanol. D) Lengths of PTs grown in vitro for 5 hours; N ≥ 203 PTs per genotype. E) Representative PTs from each genotype from (D), imaged at 10X magnification with DIC optics; scale bar represents 100 μm.

We reasoned that increased seed production in *frh3* plants could be caused by direct improvement of growth and fertility of *hpat1/3* PTs, or possibly due to improved reception of the female tissue to complement poor PT growth. To distinguish between these two modes, we performed reciprocal crosses between *hpat1/3* and *frh3* plants. *frh3* pistils crossed with *hpat1/3* pollen produced few seeds, with silique sizes and seed sets resembling those of normal *hpat1/3* plants, while *hpat1/3* pistils crossed with *frh3* pollen resulted in large, full siliques (Figure 1B). Therefore, seed production in *frh3* is specifically caused by improved fertility of *hpat1/3* pollen.

To learn more about how pollen fertility was improved in *frh3*, we first asked if PT growth was also improved. *frh3* PTs grown for 4 hours *in vitro* were significantly longer than *hpat1/3* PTs and approached near-WT lengths (Figure 1D, E). High bursting frequency and increased width of *hpat1/3* PTs were also significantly rescued in *frh3* (Figure 2A, C), suggesting that the compromised structural integrity of the cell wall caused by *hpat1/3* was suppressed. *hpat1/3* PTs also germinated at a slightly lower frequency compared to WT, and this was not rescued in *frh3* PTs (Figure 2B). Because PT germination requires secretion of cell wall materials, and we previously showed that knocking down the secretory pathway rescued *hpat1/3* PT growth defects (Beuder et al., 2020, 2022), we wanted to examine how the secretion of HPAT-modified cargos was affected in *frh3* PTs. To do this, we crossed the GF(EXT3)P reporter into *frh3* and measured the amount of fluorescent signal secreted into the cell wall (as described in Beuder et al., 2020). Apical secretion of the GF(EXT3)P reporter was higher in *hpat1/3* compared to WT PTs (Figure 2D), consistent with our previous observations (Beuder et al., 2020). This was partially rescued in *frh3*, although GF(EXT3)P secretion in *frh3* was higher than in WT (Figure 2D). In summary, our experiments suggest that improved growth and fertility of *frh3* PTs may be due to the suppression of cell wall defects associated with the loss of HPAT activity, and this may occur through reduced secretion of extensin-related cargos (and possibly other cell wall materials).

**Figure 2.**
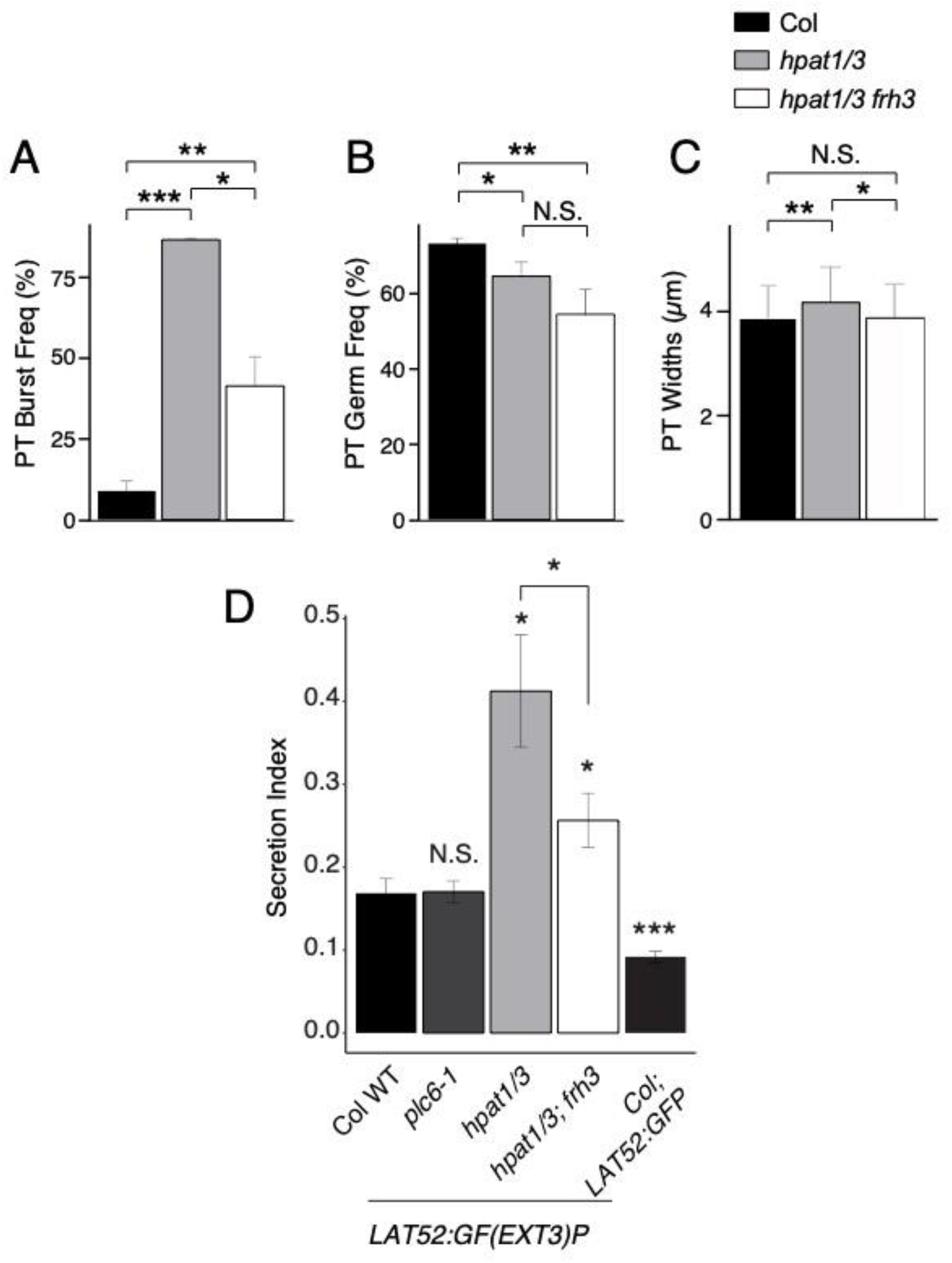
*hpat1/3* cell wall-related defects are also suppressed in *frh3* PTs. A) PT bursting and (B) germination frequencies measured after 2 hours growth *in vitro*. Experiments were repeated in triplicate for each genotype. N_bursting_ ≥ 155 PTs analyzed per experiment per genotype. N_germation_ ≥ 239 per experiment per genotype. C) PT widths measured after 1 hours, with N ≥ 101 PTs per genotype. Statistical analyses performed using Student’s t-test. D) Secretion indices for PTs expressing *LAT52:GF(EXT3)P* reporter, as well as the non-secreted control *LAT52:GFP*. Bars represent average ratios and error bars represent standard error. Statistical analyses performed using Student’s t-test compared to WT, except where noted; * represents p-value < 0.05, ** represents p-value < 0.005, *** represents p-value < 0.0005.

### 3.2 *frh3* mapped to a mutation in PLC6

To identify the *hpat1/3* suppression-causing mutation in *frh3*, we performed whole-genome sequencing of BC4F2-generation *frh3* suppressor plants, as well as non-suppressed siblings and *hpat1/3* plants. Sample preparation, sequencing strategy, bioinformatic analysis, variant filtering, and other steps were performed as previously described (Beuder & MacAlister, 2020). This led to the identification of a guanine to adenine substitution at genomic position 16754174 of chromosome 2, which mapped to *AtPLC6* (At2g40116) (Supplemental Table 2). This mutation is predicted to change glutamic acid to lysine at amino acid position 569 in exon 9, and we refer to this allele as *plc6-4* (Supplemental Figure 1). The PLC6 protein is predicted to contain a normal arrangement of domains including an EF hand domain, X and Y catalytic domains, and a lipid-binding C2 domain (Mueller-Roeber & Pical, 2002). Position 569 is near the C-terminus within the C2 domain (Supplemental Figure 1B).

If *plc6-4* is the *hpat1/3* suppressor mutation, then it should co-segregate with the suppressed phenotype as we propagate the *frh3* line. To test this, we backcrossed *frh3* plants with *hpat1/3* plants and allowed the F1s to self-fertilize. Out of 65 F2 progeny genotyped for the *plc6-4* allele, we identified only two plants that were homozygous WT for *PLC6*, indicating that there was a strong transmission bias for *plc6-4* compared to *PLC6* in the *hpat1/3* background (Supplemental Figure 2B). Additionally, the suppressed phenotype perfectly co-segregated with *plc6-4*, as the two F2 *PLC6 +/+* plants recovered in this generation were phenotypically identical to non-suppressed *hpat1/3* plants (Supplemental Figure 2C, D). Plants that were heterozygous for *PLC6 (hpat1/3; PLC6/plc6-4*) had significantly higher seed counts than those that were *PLC6^WT^ (hpat1/3; PLC6/PLC6*), indicating that seed set is markedly improved when only half of the pollen carry the suppressive mutation; however, seed counts were highest in triple homozygous mutants (*hpat1/3; plc6-4/plc6-4*) (Supplemental Figure 2C, D).

To confirm that *plc6-4* is the *hpat1/3* suppressor mutation in *frh3*, we tested the ability of the WT *PLC6* gene to rescue the suppressor phenotype (i.e. reverting the high-fertility phenotype of suppressed pollen to the low-fertility phenotype of *hpat1/3* pollen). We cloned a 3405 bp fragment that included the *PLC6* genomic sequence and upstream intergenic region (native promoter) into the binary vector pFAST-G01, which contains a convenient seed coat GFP selection marker (Shimada et al., 2010), and transformed this into *frh3* plants. Positive transformants were crossed reciprocally with WT, and we examined the resulting ratios of GFP+:GFP-seeds to measure the transmission rate of the transgene. Seven T1 plants were identified as having single-locus transgene insertions when crossed as females with WT pollen (Supplemental Figure 3). The GFP+:GFP-seed ratios were then examined from crosses in which these seven T1 plants were used as males. For all seven T1s, there was a statistically significant decrease in the number of observed GFP+ seeds compared to the expected value, indicating that the presence of the *PLC6* transgene specifically reduced pollen transmission (Supplemental Figure 3). This result is consistent with the presence of the *PLC6p:PLC6* transgene restoring normal *PLC6* activity in *frh3* pollen, counteracting the suppressive effect conferred by the *plc6-4* mutation and restoring the *hpat1/3* low fertility phenotype. Therefore, *plc6-4* was indeed the causal mutation in the *hpat1/3* suppressor family *frh3*, and *plc6-4* is a recessive mutation, consistent with a loss of function allele.

### 3.3 *plc6-4* decreases pollen transmission *in vivo* and polarized growth *in vitro*

To learn more about *PLC6’s* role in PT growth, we wanted to examine how the *plc6-4* mutation affects pollen fertility in the absence of the *hpat1/3* mutations. We outcrossed *hpat1/3: plc6-4* suppressors with WT plants, allowed the F1s to self-fertilize, and genotyped the F2s to identify *plc6-4* plants that were homozygous WT for both *HPAT1* and *HPAT3*. To compare the transmission efficiency between mutant *plc6-4* and wild-type *PLC6* pollen, *plc6-4* heterozygotes (*HPAT1/3; PLC6/plc6-4*) were outcrossed as males with WT females. We genotyped 98 progeny for the presence or absence of the *plc6-4* allele and discovered an approximate 2:1 ratio of *PLC6/PLC6: PLC6/plc6-4* progeny, which differed significantly from a 1:1 ratio expected for normal transmission (chi squared p-value = 5.93E-4) and indicates that *plc6-4* PTs were outcompeted by WT.

To identify the underlying causes of decreased *plc6-4* pollen transmission, we performed *in vitro* PT growth assays. Interestingly, *plc6-4* PTs appeared drastically different compared to WT, with curvy and hooked shapes signifying erratic directional changes during growth (Figure 3A, and quantified in Figure 7C). This phenotype was not observed in *hpat1/3; plc6-4* PTs, whose appearance strongly resembled WT (Figure 1E), indicating that *hpat1/3* suppresses the abnormal “crooked growth” phenotype of *plc6-4* PTs. We also observed decreased length and increased width in *plc6-4* PTs, indicating that cell polarity during tip growth is compromised (Figures 3B, D). *plc6-4* PTs also displayed lower germination frequencies, a phenotype shared with the exocyst secretion mutants previously characterized (Figure 3D; Beuder et al., 2020). *plc6-4* self-fertilized siliques contained fewer seeds than WT (Figure 3C, D). Notably, positions with no seeds were typically found near the basal end of the silique (Figure 3C), suggesting that decreased PT elongation is a primary cause of lower seed counts.

**Figure 3.**
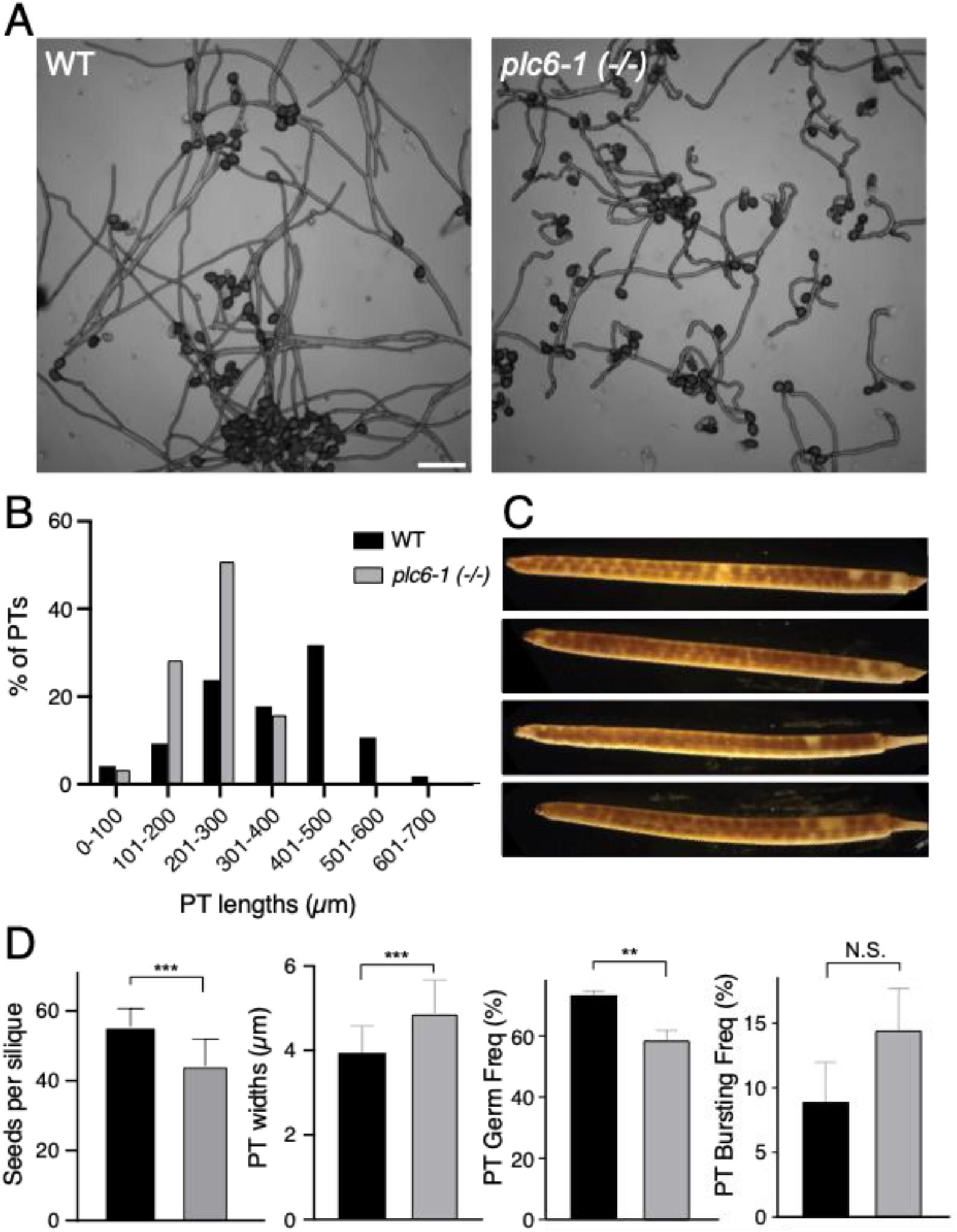
*plc6-4* PTs are curlier, wider, and germinate less frequently than WT. A) PTs grown in vitro for four hours and imaged with DIC optics at 10X magnification; scale bar represents 100 μm. B) Quantification of PT lengths after 5 hours growth in vitro N ≤ 202 PTs per genotype. C) Pictures of *plc6-4* siliques. D) Reproductive phenotypes. Far left-seed counts for WT (N = 14) and plc6-4 (N= 13). Left middle-PT width measurements after 1 hours of growth; N ≥ 200 PTs per genotype. Right middle and far right-Germination and bursting frequencies after 2 hours of in vitro growth. Both bursting and germination experiments were repeated in triplicate for each genotype. N_bursting_ ≥ 159 PTs analyzed per experiment, per genotype. N_germination_ ≥ 404 pollen grains analyzed per experiment per genotype. Statistical analyses performed using Student’s t-test; ** represents p-value < 0.005.

To examine *plc6-4* PT growth *in vivo*, we manually pollinated WT pistils with either WT or *plc6-4* pollen and collected them 24 hours after pollination. To visualize PTs within the pistil we stained the fixed pistils with aniline blue fluorochrome, which binds (1→3)-β-glucans (callose) enriched in PT cell walls. Both WT and *plc6-4* PTs appeared to grow normally through the transmitting tract and during ovule targeting (Supplemental Figure 4), suggesting that the meandering growth is suppressed *in vivo*. This could be due to the physical support provided by the pistil tissue *in vivo*, which may constrain PT grow and mask the crooked growth phenotype observed *in vitro*. Also, guidance cues provided by the pistil may be reinforcing PT growth direction, and the absence of these cues *in vitro* may cause frequent reorientation of PT growth. Taken together, this data suggests that *PLC6* promotes pollen growth by helping PTs maintain cell polarity during growth *in vitro*, but this may not be essential for *in vivo* PT growth and seed production. However, the reduced transmission of *plc6-4* pollen and the reduced seed set observed, particularly in the basal portion of *plc6-4* pistils, suggests that *PLC6* is required for full pollen fertility *in vivo*.

### 3.4 *plc6* insertion mutants do not suppress *hpat1/3* fertility defects

Because the suppressive effects of *plc6-4* were rescued by the WT *PLC6* gene, we hypothesized that *plc6-4* was a loss of function allele. Therefore, we wanted to determine if knocking out *PLC6* expression would also suppress *hpat1/3* pollen fertility defects. We obtained three *plc6* mutant lines with T-DNA insertions located in different regions of the *PLC6* gene (Supplemental Figure 1A) and checked for expression through PCR using cDNA prepared from flower RNA as the template. Using two primer sets (Supplemental Table 1), we attempted to amplify different regions (1 and 2) of the predicted transcript from WT and insertion mutant cDNA. Region 1 spans most of the catalytic X domain and the EF domain (which is also required for catalytic activity in PLCs; (Otterhag et al., 2001), and region 2 spans the second half of the catalytic Y and C2 domains (Supplemental Figure 1A, B). Both regions were amplified successfully from WT cDNA. Region 1 expression was absent from CSHL_GT4996 (*plc6-2*) and SALK_090508 (*plc6-3*) (although small amounts of aberrantly sized products were detected), while SALK_0401365 (*plc6-1*) showed that region 1 was expressed (Supplemental Figure 1D). Conversely, region 2 transcript was detected in *plc6-2* and *plc6-3* cDNA, but not *plc6-1*. We reasoned that loss of region 1 was likely to produce nonfunctional PLC6 protein products, whereas the absence of region 2 in *plc6-1* cDNA may not abolish PLC6 activity if the truncated *PLC6-1* mRNA is translated with the EF, X and half of the Y domain intact. Therefore, we decided to focus on characterizing *plc6-2* and *plc6-3*.

To test for *hpat1/3* suppression, we crossed the *plc6-2* and *plc6-3* T-DNA lines into *hpat1/3* and obtained plants that were heterozygous for either T-DNA mutation (*hpat1/3; PLC6/plc6-2* and *hpat1/3; PLC6/plc6-3*). We crossed *hpat1/3; plc6 +/-* plants as males with WT females with the expectation if either *plc6* T-DNA allele was suppressing *hpat1/3* pollen fertility defects, then we would observe a transmission bias in favor of the mutation as we found for *plc6-4*. Interestingly, we observed a significant bias against transmission for *plc6-2* and *plc6-3* (N_*PLC6 vs plc6-2*_ = 51,82.4% transmission rate of *PLC6*, chi-squared p-value = 3.82E-06; N_*PLC6 vs plc6-3*_ = 54, 74.1% transmission rate of *PLC6*, chi-squared p-value = 4.03E-4), indicating that these mutations decreased *hpat1/3* pollen fertility. We obtained *hpat1/3 plc6-2* and *hpat1/3 plc6-3* triple mutant plants and performed seed counts. Compared to their *hpat1/3 PLC6^WT^* siblings, *hpat1/3; PLC6/plc6-2* and *hpat1/3; plc6-2* triple homozygous mutants had significantly reduced seed counts (Figure 4D), as did *hpat1/3; plc6-3* triple homozygous mutants (Figure 4J). Since this is the opposite of what we expected, we examined pollen phenotypes of these mutants more closely through *in vitro* PT growth assays. PT lengths were comparable between both *hpat1/3; plc6* transgene insertion triple mutants and *hpat1/3* double mutants (Figure 4A, C, G and I). PT bursting occurred at a high frequency in *hpat1/3*, and this appeared unchanged in both triple mutants (Figure 4E, K); however, PT germination frequencies were significantly decreased in both triple mutants (Figure 4F, L). This data indicates that neither *plc6-2* nor *plc6-3* suppress *hpat1/3* pollen fertility defects; on the contrary, they acted as enhancers of the low pollen fertility phenotype. This data also suggests that the missense *plc6-4* allele is not acting as a full loss-of-function allele.

**Figure 4.**
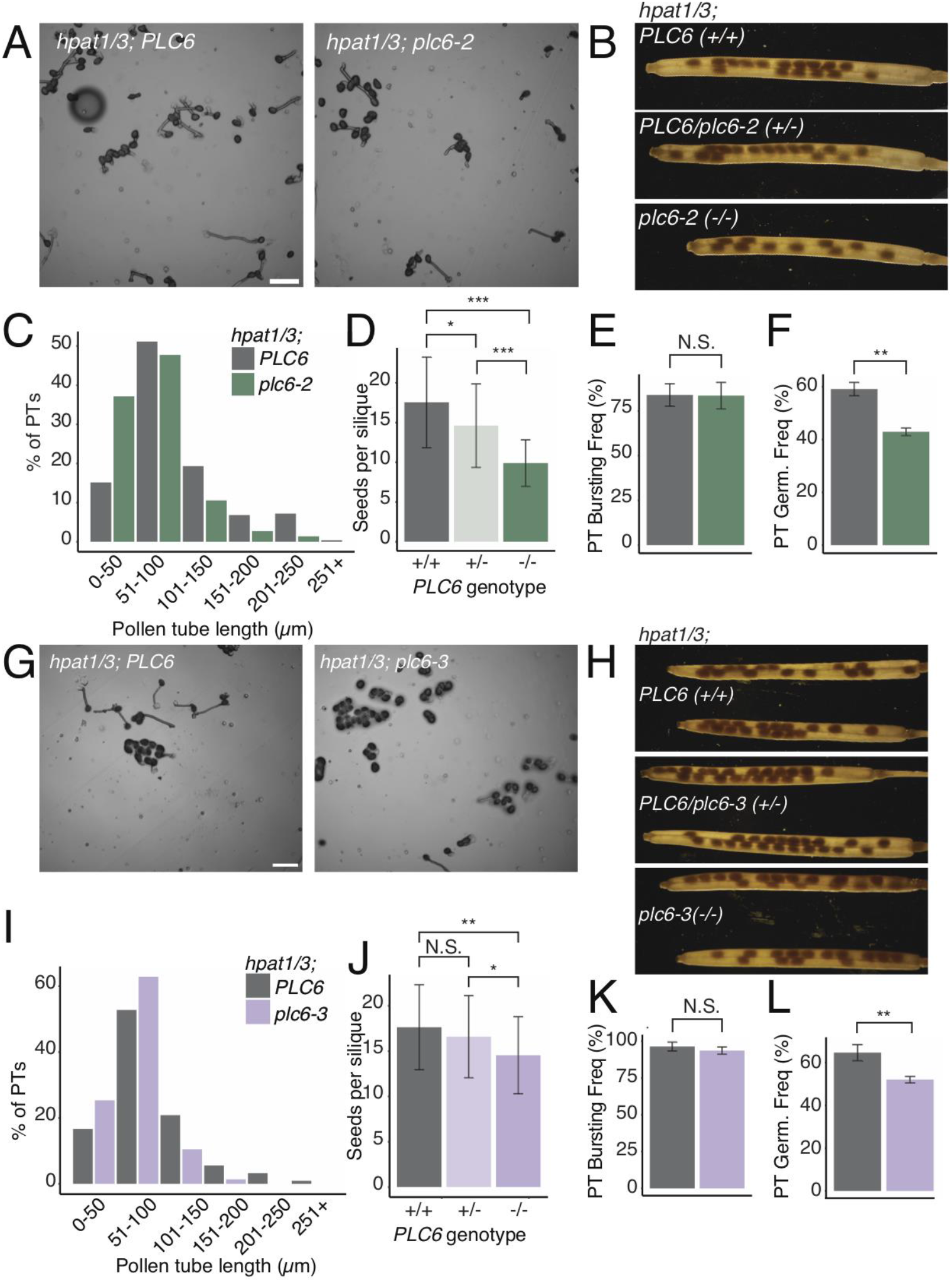
*plc6* T-DNA insertion mutations do not improve *hpat1/3* pollen fertility. A) PTs grown in vitro for 2 hours and imaged with DIC optics at 10X magnification; scale bar represents 100 μm. B) Representative siliques for each genotype cleared with 70% ethanol. C) Quantification of PT lengths after 2 hours growth in vitro N ≥ 218 PTs per genotype. D) Seed counts based on plant genotype; (-) denotes *plc6-2* mutant allele. N ≥ 19 siliques per genotype. Bars represent average seed counts and error bars represent standard deviation. E) PT bursting frequencies after 2 hours in vitro growth. Experiment repeated three times with N ≥ 112 PTs per genotype per experiment. F) PT germination frequencies after 2 hours *in vitro* growth. Experiment repeated three times with N ≥ 332 pollen grains measured per genotype per experiment. G) PTs grown *in vitro* for 2 hours and imaged with DIC optics at 10X magnification; scale bar represents 100 μm. H) Representative siliques for each genotype cleared with 70% ethanol. I) Quantification of PT lengths after 2 hours growth in vitro N ≥ 216 PTs per genotype. J) Seed counts based on plant genotype; (-) denotes *plc6-3* mutant allele. N ≥ 22 siliques per genotype. K) PT bursting frequencies after 2 hours *in vitro* growth. Experiments repeated twice for each genotype with N ≥ 150 pollen grains examined, except in one experiment, the N_*hpat1/3*_=66. L) PT germination frequencies after 2 hours *in vitro* growth. Experiment repeated three times for *hpat1/3* (N ≥ 239 per experiment) and twice for *hpat1/3; plc6-3* (N ≥ 501 per experiment). Statistical analyses performed using Student’s t-test.

To further examine the role of *PLC6* in pollen fertility, we also wanted to determine if *plc6-2* and *plc6-3* mutations affected pollen transmission and growth in an otherwise WT genetic background. We performed test crosses with plants that were heterozygous for either *plc6-2* or *plc6-3* as males (*PLC6/plc6-_*), and, surprisingly, we observed no transmission bias either for or against either transgene (N_*PLC6 vs plc6-2*_ = 79, 46.8% transmission rate of *PLC6*, chi-squared p-value = 0.57; N_*PLC6 vs plc6-3*_ = 44, 47.6% transmission rate of *PLC6*, chi-squared p-value = 0.75). We compared seed sets of *plc6-2* and *plc6-3* homozygous mutants to WT and found that the seed counts appeared similar to WT (Figure 5B, D, F and H). Furthermore, the lengths and overall appearance of *in vitro*-grown *plc6-2* and *plc6-3* PTs both strongly resembled WT (Figure 5A, C, E and G). In summary, the *plc6-2* and *plc6-3* mutations decreased fertility and *in vitro* germination in *hpat1/3* PTs, but these mutations did not cause observable effects in the WT background.

**Figure 5.**
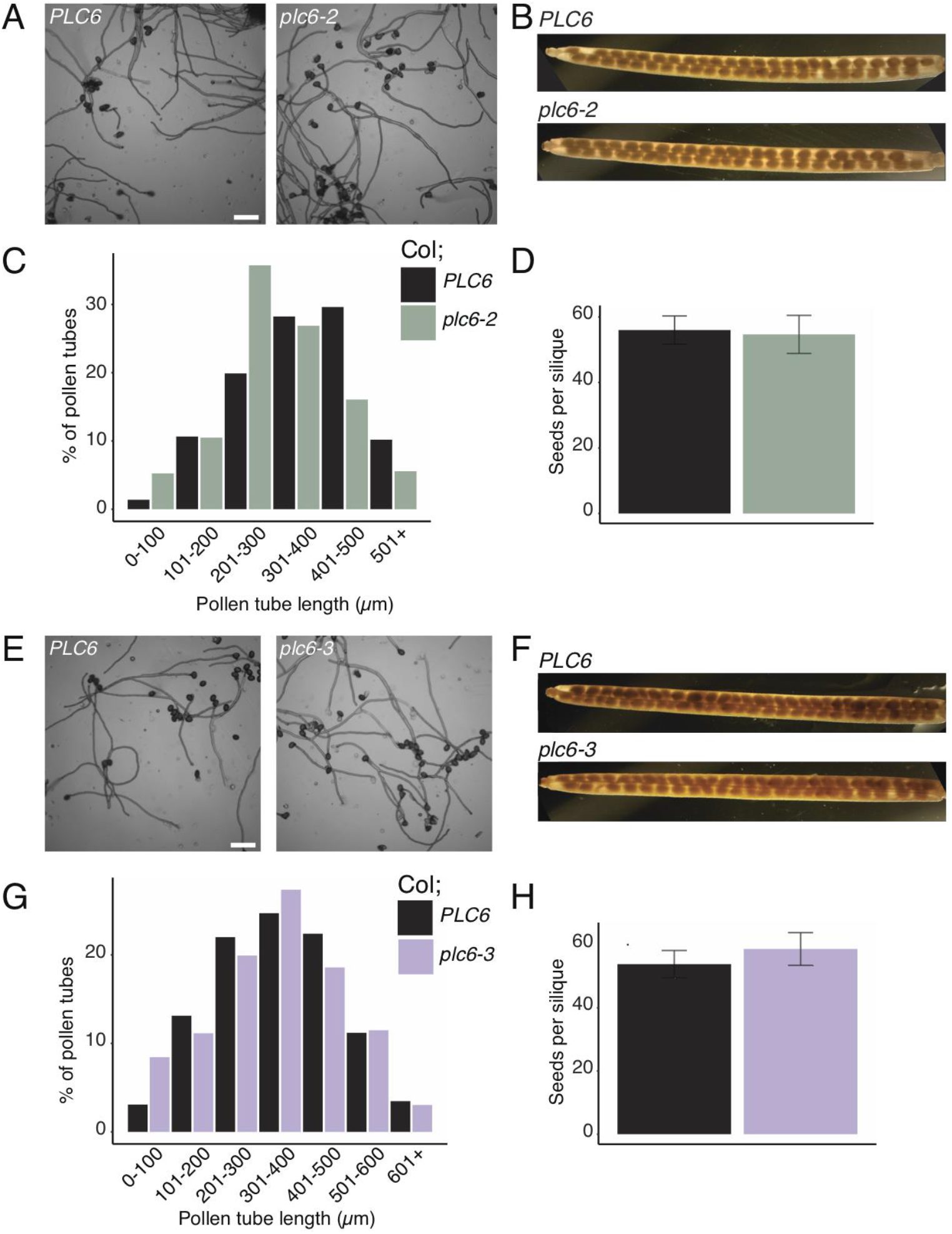
Seed production and in vitro PT growth are not disrupted in *plc6* T-DNA insertion mutants. A-D) Reproductive phenotypes in WT vs *plc6-2* mutant. A) PTs grown *in vitro* for 4 hours and imaged with DIC optics at 10X magnification; scale bar represents 100 μm. B) Representative siliques for each genotype cleared with 70% ethanol. C) Quantification of PT lengths after 4 hours growth in vitro. N ≥ 216 PTs per genotype. D) Seed counts for each genotype; (-) denotes *plc6-2* mutant allele. N ≥ 19 siliques per genotype. E-H) Reproductive phenotypes in WT vs *plc6-3* mutant. E) PTs grown *in vitro* for 4 hours and imaged with DIC optics at 10X magnification; scale bar represents 100 μm. F) Representative siliques for each genotype cleared with 70% ethanol. G) Quantification of PT lengths after 4 hours growth *in vitro*. N ≥ 259 PTs per genotype. H) Seed counts for each genotype; (-) denotes *plc6-3* mutant allele. N ≥ 19 siliques per genotype. Statistical analyses performed using Student’s t-test.

### 3.5 *PLC* gene expression across reproductive and vegetative tissues

A possible explanation for why we did not observe an abnormal PT growth phenotype in either *plc6* T-DNA insertion mutants could be functional redundancy among pollen-expressed *PLCs*. To explore this possibility, we wanted to compare individual *PLC* gene expression in pollen. There are 9 *PLC* genes in the Arabidopsis genome. *PLC1, 2, 3*, and *7* are most closely related to *NtPLC3* and Petunia (Pet) *PLC1* (Figure 6A), which have important roles in PT growth (Dowd et al., 2006a; Helling et al., 2006).

**Figure 6.**
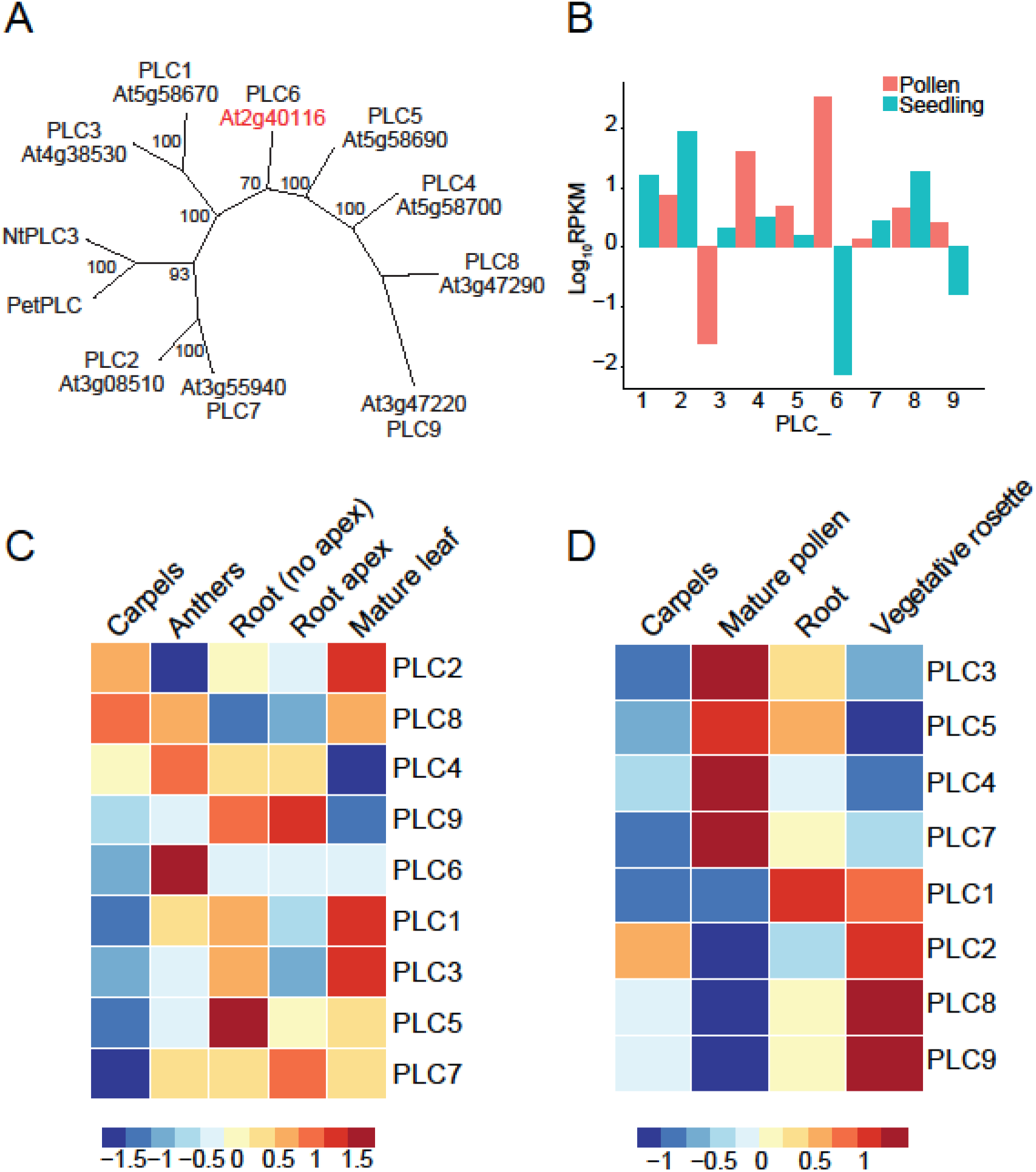
PLC phylogenetic analysis and expression profiles in Arabidopsis. A) Phylogenetic analysis of PLCs from Arabidopsis, Petunia (Pet), and tobacco (Nt). Unrooted maximum likelihood tree shown with bootstrap values for each node. B) RNA sequencing analysis showing RPKM values transformed to log_10_ scale for each *PLC* gene in *Arabidopsis* pollen or seedling samples. Dataset obtained from Lorraine et al. 2013. C) Relative expression on log_10_ scale of each *PLC* gene. Heatmap generated with RNA sequencing data from Kleplikova Atlas (Klepikova et al. 2016). D) Relative expression on log10 scale of each PLC gene. Heatmap generated with microarray data from Schmid et al. 2005.

We plotted *PLC* gene expression levels using publicly available RNA-sequencing data for pollen and seedlings (Loraine et al., 2013) (Figure 6B). Notably, *PLC6* was found to be the most highly expressed *PLC* in pollen, with an RPKM value almost one order of magnitude higher than the second most-highly expressed *PLC* in pollen, *PLC4*. In addition to *PLC4* and *6, PLC5* and *9* also had higher levels of expression in pollen than in seedlings. Similarly, RNA-sequencing data published on the Klepikova Atlas (Klepikova et al., 2016), showed that *PLC6* expression was highly enriched in the anthers (pollen was unavailable). *PLC6* expression was also highest in anthers among all PLCs, followed by *PLC4* (Figure 6C). We also generated expression profiles using microarray data (Schmid et al., 2005) for each *PLC* gene except for *PLC6*, for which data was unavailable, and observed that *PLC3, 4, 5*, and *7* expression was highly enriched in mature pollen (Figure 6D).

These analyses indicate that *PLC6* is highly expressed and enriched in pollen and anther, and its expression profile is unique among the *PLCs*, suggesting that it is a major PLC isoform present in growing PTs. However, other *PLCs* are expressed in pollen and may be important for pollen fertility.

### 3.6 The *plc6-4* mutation occurs in the C2 domain at a calcium-coordinating motif

Because our data suggests that *plc6-4* is not a full loss of function allele, we wanted to learn more about how the *plc6-4* mutation may affect PLC6 protein function by further examining the nature of the mutation. The E569 position is located in the C2 domain, and this residue is conserved in five out of nine Arabidopsis *PLC* sequences (Supplemental Figure 1C). We also aligned the primary structure of PLC6’s C2 domain to the C2 domains of three protein kinase Cs from rat and human, for which protein structures have already been determined (Corbalan-Garcia & Gómez-Fernández, 2014; Pike et al., 2007; Sutton & Sprang, 1998; Verdaguer et al., 1999). This revealed that the E569 position in PLC6 falls between key aspartic acid residues involved in calcium-coordinating, as well as other residues involved in PI interactions (Fig S5A). To determine the spatial positioning of this residue we used a homology modeling approach using the Phyre 2 protein fold recognition server (Kelley et al., 2015). The full-length, predicted PLC6 protein sequence was modeled against a protein fold library and the best model returned was fold library ID c1djyB, which was modeled using a crystal structure for the PI-specific PLC C δ1 from rat complexed with inositol-2,4,5-trisphosphate (Essen et al., 1997) as a template; this model returned a with a 100% confidence score for 87% of the PLC6 sequence. The mutated residue E569 occurred on an external loop in the C2 domain (Supplemental Figure 5B), and the charge inversion (E569K) of the *plc6-4* allele is likely to interfere with the electrostatic interactions taking place in this region, such as calcium binding.

Based on our *in silico* analysis of the *PLC6* C2 domain, we hypothesized that the *plc6-4* mutation may be inhibiting PLC6’s ability to bind and coordinate calcium. To test this, we examined *plc6-4* PT growth response across different calcium levels. We performed *in vitro* growth assays to analyze both PT elongation and degree of meandering in response to varying calcium concentrations in the growth media. After 2.5 hours of growth on standard (2 mM) calcium, *plc6-4* PTs were shorter than WT, as we had observed previously (Figure 7A, B). WT PTs were largely insensitive to changes in calcium level with a statistically significant reduction in average length observed only at 8 mM calcium levels compared to 2 mM; however, *plc6-4* PT elongation was optimal at 1 mM calcium, and decreased as calcium levels increased (Supplemental Table 3).

**Figure 7.**
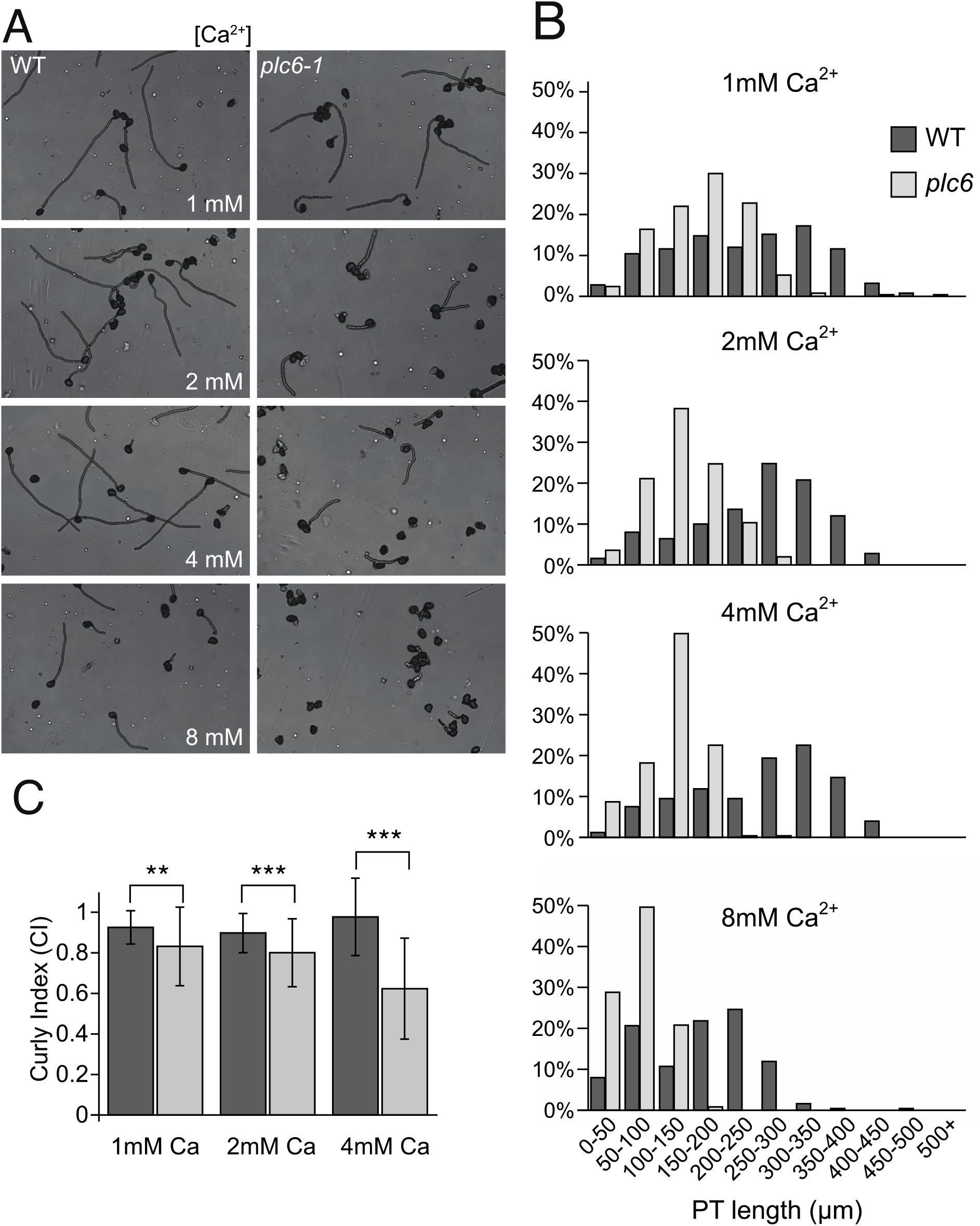
*plc6-4* PT growth *in vitro* is inhibited by increased calcium levels. A) PTs grown for 2.5 hours and imaged with DIC optics at 10X magnification; B) PT lengths quantified after 2.5 hours, with N ≥ 250 PTs per experiment per genotype. C) Length/distance ratios to measure PT straightness. Measurements taken after 2.5 hours of growth. N ≥ 64 PTs per genotype per experiment. Statistical analyses performed using Student’s t-test

The meandering growth of PTs was measured with a “straightness index (SI)”, the ratio of the total length of the pollen tube divided by the shortest distance between the pollen grain and the pollen tube tip. An SI value of 1 indicates that the PT was perfectly straight, and a lower SI indicates meandering growth. At each tested calcium concentration, the WT PTs were significantly straighter than *plc6-4* (Figure 7C). In both genotypes, straightness was unaffected in 1 mM calcium compared to 2 mM (Table S4). *plc6-4* PTs were less straight at 4 mM than at 2mM (Supplemental Table 4), while WT PTs were actually straighter at 4 mM than at 2 mM. This data indicates that in addition to elongation, *plc6-4* PT straightness is also negatively affected by higher calcium levels.

### 3.8 Subcellular localization of PLC6-mNG

Previous research shows that PI(4,5)P_2_ availability is a limiting factor for EXO70 binding at the plasma membrane (Pleskot et al., 2015). PI5P kinase overexpression results in PI(4,5)P_2_ accumulation, meandering PT growth and decreased PT elongation (Ischebeck et al., 2008; Zhao et al., 2010). Therefore, it is possible that the PLC6-4 protein may be defective in PI(4,5)P_2_ binding and/or cleavage to form IP3 and DAG, which would expand PI(4,5)P_2_ availability at sub-apical regions along the PT plasma membrane and cause dispersion of exocyst-mediated secretion; this would predictably decrease cell polarity and tip growth. In this model, apical accumulation of secreted cargos would decrease at the tip and restore extensibility and polarized growth in *hpat1/3* PTs, but in the *plc6-4* single mutant background, loss of polarized exocytosis misdirects apical secretion, leading to curly growth paths and wider PTs. Therefore, we wanted to examine how PLC6 accumulated at the apical plasma membrane.

To determine the subcellular localization of PLC6, we cloned and transformed plants with a construct encoding the *PLC6* genomic sequence fused with a C-terminal mNeonGreen (mNG) under the native *PLC6* promoter (*PLC6p:PLC6-mNG*). PI(4,5)P_2_ and other PLCs have been shown to localize to the apical and subapical plasma membrane in PTs, respectively, (Dowd et al., 2006b; Helling et al., 2006) so we hypothesized that PLC6-mNG would localize to the subapical plasma membrane as well. Using confocal microscopy, we examined the pattern of mNG signal in PTs from three positive, independent T1 plants. We observed diffuse signal throughout the cytoplasm, with no enrichment at the plasma membrane (Figure 8). However, to our surprise, we observed enrichment of mNG signal in the shape of two connected circles far from the apical tip. Using DAPI to visualize the sperm and vegetative nuclei, we confirmed that mNG signal was enriched in the endomembrane surrounding the sperm nuclei (Figure 8). These patterns were consistent among all three T1 plants.

**Figure 8.**
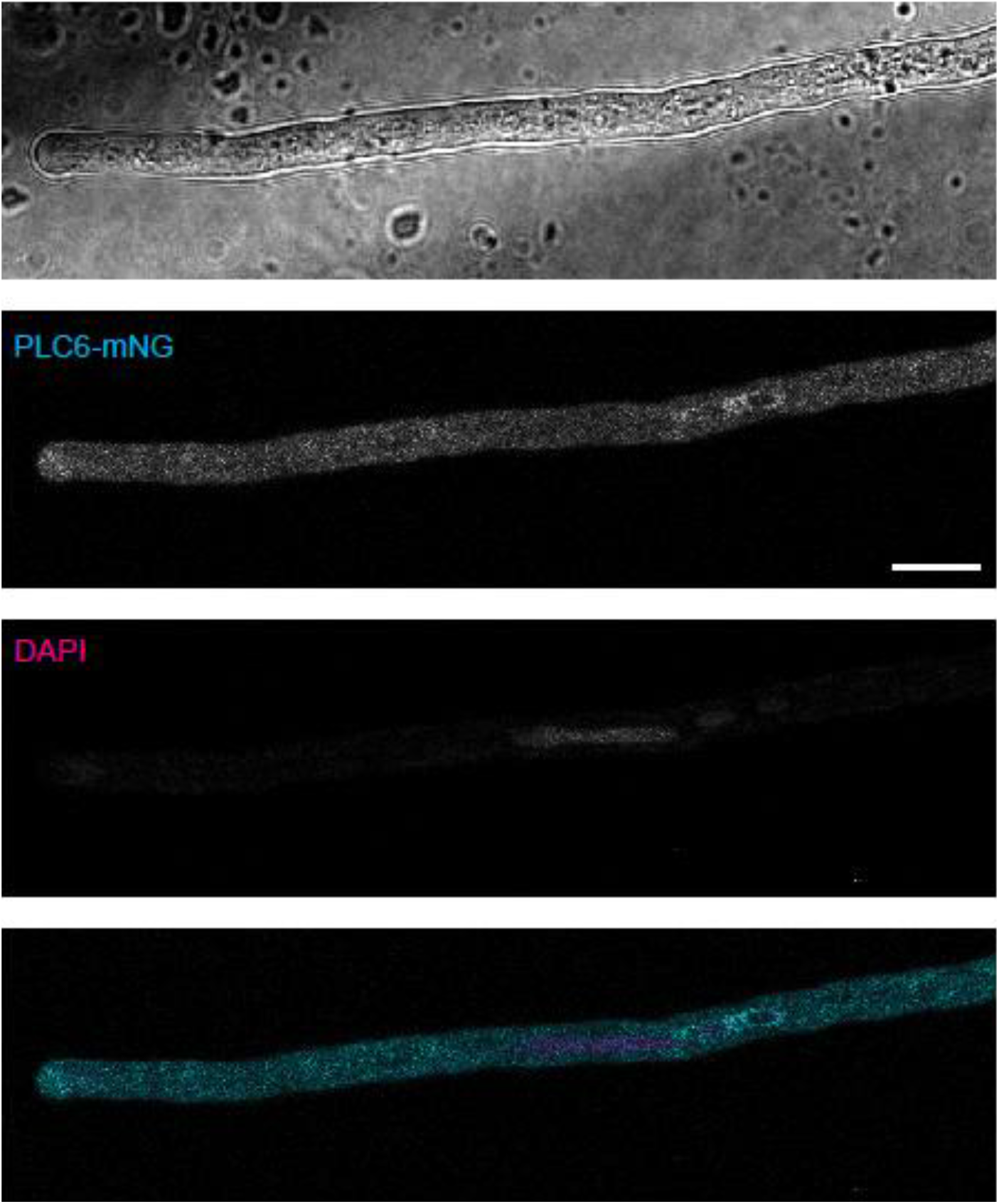
*PLCp:PLC6-mNG* is expressed in PTs and is enriched around sperm nuclei. Medial z-slice of a representative PT expressing *PLC6p:PLC6-mNG*, imaged with confocal microscopy at 100X magnification. In merged image, mNG is false-colored cyan and DAPI is false-colored magenta. Scale bar represents 10 μm.

To determine if the PLC6-mNG C-terminal fusion protein was functional, we wanted to test its ability to rescue suppression of *hpat1/3; plc6-4* pollen in the same manner as described earlier (Supplemental Figure 3). We transformed the construct into *frh3* suppressor plants and crossed positive T1s reciprocally with WT. For eight single loci-insertion T1 plants, four of these plants produced significantly less than 50% fluorescent seeds when used as pollinators, indicating that suppression was being rescued in *hpat1/3; plc6-4* pollen carrying the construct, which suggests the PLC6-mNG fusion protein is functional (Supplemental Figure 6). The other four T1 plants produced seed sets with approximately a 1:1 ratio of fluorescent to non-fluorescent seeds, indicating no transmission bias for pollen carrying the construct. We believe this data suggests that the PLC6-mNG C-terminal fusion protein is likely functional and it is possible that the four T1 plants that did not show a bias could be due to low expression of PLC6-mNG; however, we cannot be certain that mNG is not affecting PLC6 localization and/or function.

## 4. Discussion

Here, we report that a missense mutation in *PLC6*, which corresponds to an E569K amino acid substitution, strongly rescued poor pollen fertility of *hpat1/3* by improving PT elongation, decreasing morphological defects, and decreasing bursting frequency. The elevated apical secretion of the GF(EXT3)P reporter protein in *hpat1/3* is decreased in *plc6-4* PTs (Figure 2D), suggesting that *PLC6* also plays a role in promoting secretion of Hyp *O-*arabinoside-containing cargos. *plc6-4* also decreased pollen germination, as does *plc6-2 and plc6-3* (in the *hpat1/3* background), consistent with a role of *PLC6* in promoting secretion. Because these phenotypes are similar to what we previously observed in the *hpat1/3;exo70a2-2* and other secretion mutant suppressors (Beuder et al., 2020, 2022), and PLCs are known to regulate PI(4,5)P_2_ availability at the plasma membrane, we reasoned that *PLC6* may influence secretion by regulating exocyst binding site availability and affecting apical secretion; however, there are important phenotypic differences to note between the *plc6-4* and the other known *hpat1/3* suppressor mutations.

In the *plc6-4* single mutant background, GF(EXT3)P secretion was not affected, but it was decreased in *exo70a2* and *sec1a* mutants (Figure 2D, Beuder et al., 2020, 2022). Compared to *plc6-4*, PT germination is decreased to a larger extent in *exo70a2* and *sec15a* PTs (Figure 3D; (Beuder et al., 2020). While *exo70a2* and *sec15a* PTs did not elongate as much as those of WT, they were phenotypically normal otherwise (Beuder et al., 2020), which contrasts to the looped and crooked appearance of *plc6-4* PTs. Because of the phenotypic differences between *plc6-4* and exocyst mutant PTs, *plc6-4* is not likely causing robust knockdown of global exocyst-mediated secretion in the same manner as *exo70a2* or *sec15a*. This was further supported by the observation that PLC6-mNG signal was not localized at the plasma membrane, but rather to the cytoplasm, and was enriched in the sperm endomembrane system.

The ER is considered the main internal calcium store in animals (Segal et al. 2014), but other organelles likely contribute to cytoplasmic calcium levels. In human sperm, the nuclear envelope and surrounding endomembrane compartments contain calcium channels, which may be involved in regulating cytoplasmic or nuclear calcium levels (Costello et al., 2009). Less is known about calcium storage in plant cells, but cyclic nucleotide-gated ion channels (CNGCs) that transport calcium are localized to the nucleus (Costa et al., 2018). Glutamate receptor-like channels (GLRs) transport calcium through the plasma membrane to modulate cytoplasmic calcium concentrations and control PT growth, and in *Arabidopsis thaliana (At), glr1* knockout mutant PTs are less fertile due to defects in tip growth including slower growth rates and a crooked appearance, similar to *plc6-4* PTs (Michard et al., 2011). Internal calcium oscillations are also disturbed in *glr1* PTs (Michard et al., 2011). AtGLR3-GFP localizes to the sperm endomembrane in PTs (Wudick et al., 2018), suggesting that this organelle may have a role in storing calcium to regulate cytoplasmic or nuclear calcium concentration. Therefore, it is possible that rather than regulating PI(4,5)P_2_ at the plasma membrane, PLC6 may control PT growth by locally regulating PI signaling at the sperm endomembrane, which may affect calcium transport. However, to our knowledge, PI(4,5)P_2_ abundance at the PT sperm endomembrane has not been clearly demonstrated. How the *plc6-4* mutation affects PLC6 lipase activity and internal calcium oscillations are important questions to address in future research.

The transgenic construct containing the WT *PLC4* genomic sequence rescued suppression (i.e. decreased transmission) of *hpat1/3;plc6-4* pollen, which suggests that *plc6-4* is a recessive mutation, which is consistent with a loss of function allele; however, the *plc6-2* and *plc6-3* T-DNA insertion alleles did not suppress *hpat1/3* PT defects (in fact, they decreased *hpat1/3* PT fertility), indicating that *plc6-4* is likely not acting as a true null allele. Furthermore, because the *plc6-2* and *plc6-3* T-DNA insertion alleles did not cause any negative effects on WT PT growth and/or fertility, we conclude that *PLC6* is not ultimately required for these processes; but what about other PLCs? Our expression analyses indicate that *PLC6* is most highly expressed in pollen, but other *PLCs* are also expressed. One possibility for why there is no abnormal pollen phenotype observed in the *plc6-2* and *plc6-3* T-DNA lines (in the absence of *hpat1/3*) could be due to functional redundancy among the other pollen-expressed *PLCs*. Because *PLC4* expression was consistently enriched in male tissue across all datasets, it may be compensating for the absence of *PLC6* in pollen, but participation of other *PLCs* cannot be ruled out. It is also possible that PLCs have little, if any, influence on PT growth in WT *Arabidopsis. hpat1/3* PTs are already highly compromised due to defects in the structural integrity of the cell wall, so they may be more sensitive to changes that would have no effect otherwise. Analyzing PT phenotypes of higher-order *plc* mutants will determine if PLC activity, and which *PLCs*, are important for PT growth and fertility.

## 5. Acknowledgments

This material is based upon work supported by the National Science Foundation under Grant No. IOS-1755482. We thank Gregg Sobocinski for microscopy assistance and advice. Steven Beuder also received funding from the University of Michigan through the Elizabeth Youngman Fellowship (Rackham Graduate School) and the PhD Candidate Student Research Grant (Department of Molecular, Cellular, and Developmental Biology).

## 6. Author Contributions

Cora MacAlister designed the study and conducted the *frh* mutant screen. Steven Beuder wrote the paper and carried out microscopy, bioinformatic analysis of whole genome sequences to identify candidate suppressor mutations, functional rescue assays, and performed crosses and PT phenotyping. Xuesi Hua performed genotyping and PT phenotyping. Alexandria Dorchak carried out the cloning.

---

**Supplemental Figure 1.**
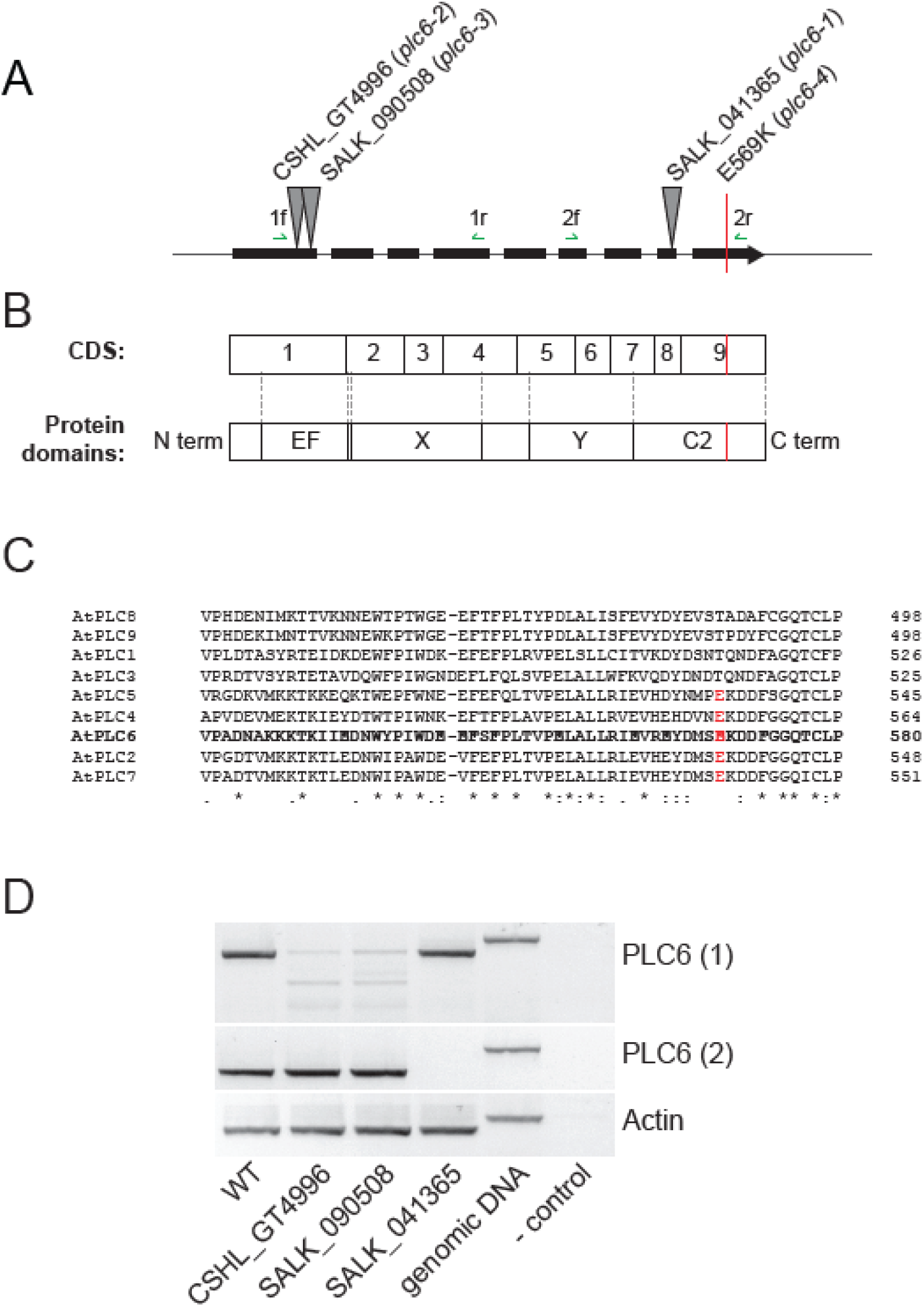
PLC6 gene map and genotyping *plc6* mutants. A) Map of the PLC6 gene indicating exons (boxes), introns, and locations of transgene insertions for *plc6-1, plc6-2* and *plc6-3*, as well as the location of the *plc6-4* mutation. Also shown are primer combinations to detect the *PLC6* transcript. B) Map of CDS corresponding to protein domains, (adapted from Cheng et al., 2017; Mueller-Roeber & Pical, 2002). C) Partial protein sequences for Arabidopsis PLCs with the E569 position indicated in red. Alignment performed with CLUSTAL omega. “*” indicates perfect alignment, “:” indicates site among group with strong similarity, and “.” Indicates site among group with weak similarity. D) Gel showing PCR products amplified from cDNA using primer combinations 1 (1F + 1R) and 2 (2F + 2R) (Supplemental Table 1), and actin. cDNA libraries were prepared from flower mRNA.

**Supplemental Figure 2.**
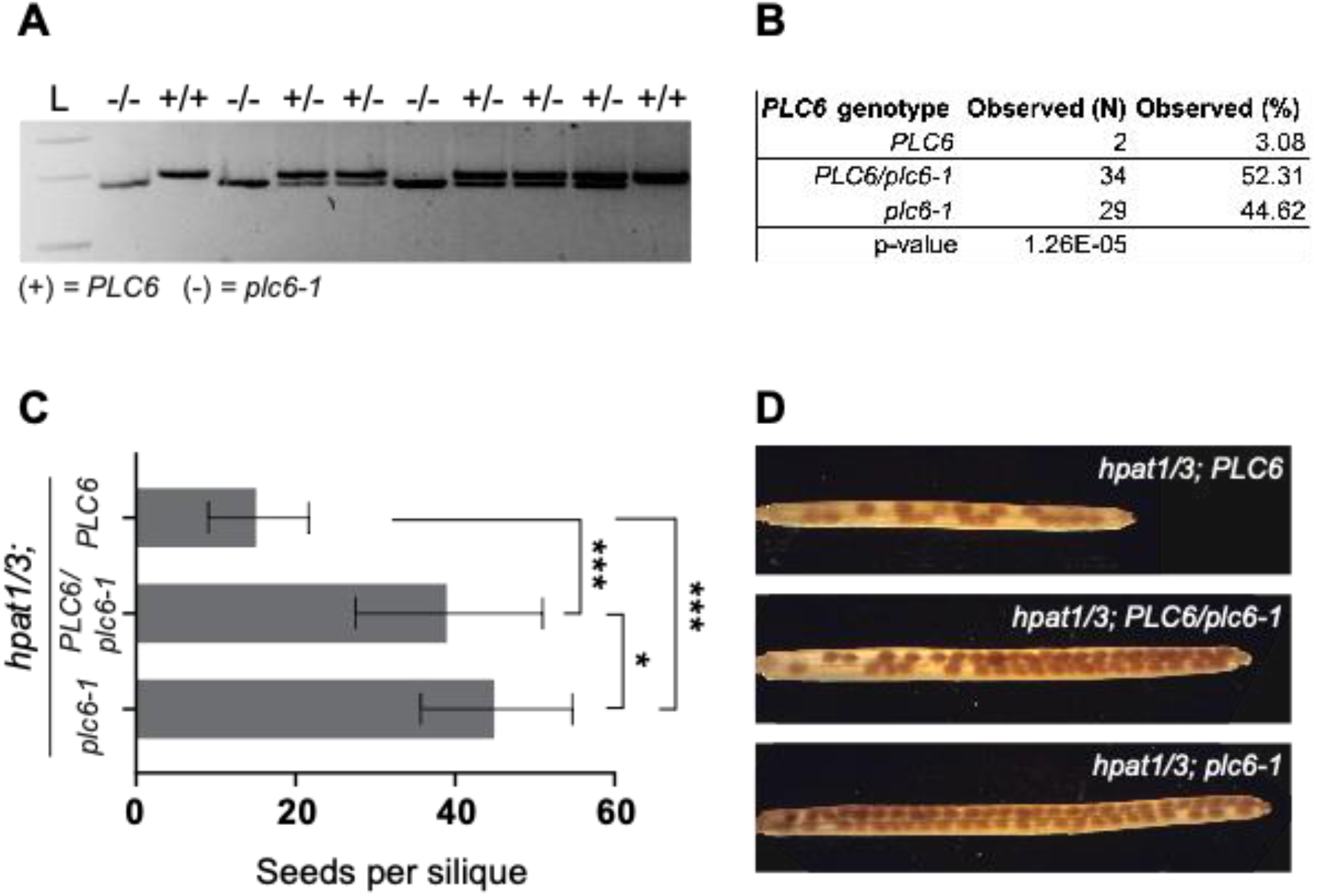
*hpat1/3* suppression phenotype co-segregates with *plc6-4* mutation in BC5F2 generation *frh3* plants. A) Genotyping of *plc6-4* mutation. PCR products were amplified using dCAPS primers, treated with HYP188III restriction enzyme, and run on a 2% agarose gel. Full-length (WT) PCR product = 207 bp. PCR product containing the *plc6-4* mutation is cleaved at the 3’ end of primer resulting in 183 and 24 bp bands. B) Genotype segregation ratios of BC5F2 individuals. C) Seed counts for BC5F2s based on PLC6 genotype. (+) denotes the WT allele, and (-) denotes *plc6-4* mutant allele. Statistics performed using Student’s t-test; * indicates p-value < 0.05. D) Representative siliques of each genotype cleared with 70% ethanol.

**Supplemental Figure 3.**
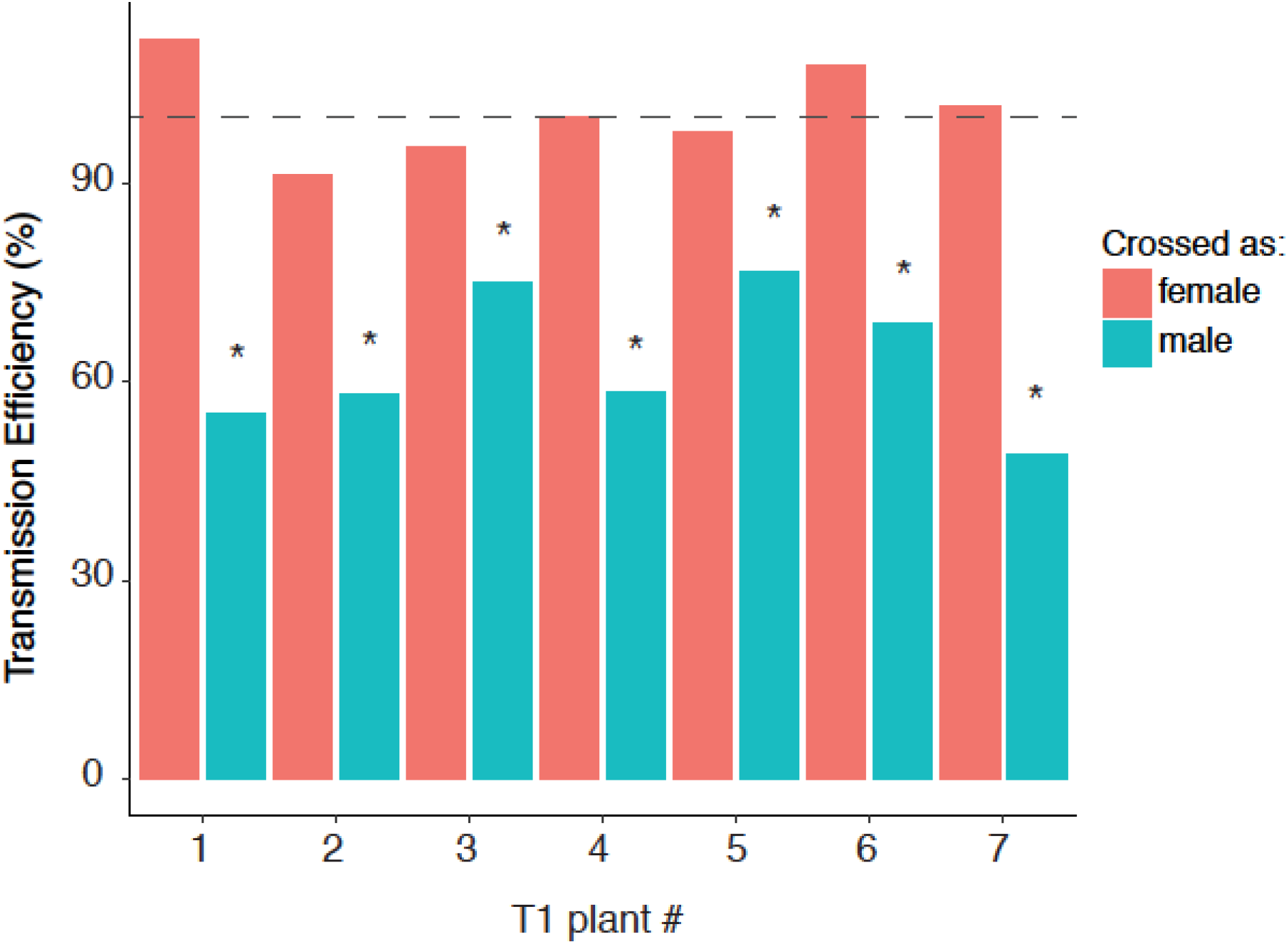
Transmission efficiency is decreased in *frh3* PTs carrying PLC6 genomic transgene. The PLC6promoter:PLC6genomic sequence was cloned into pFAST-G01 (Shimada et al. 2010) and transformed into *frh3* plants, which includes the OLE1:GFP selection marker and the genomic PLC6 sequence. Transmission rates calculated as percent of GFP+ seeds produced from a cross using transformed plants (T1-T7), and scaled to 100% by multiplying by 2 (50% GFP+ seeds means 100% transmission rate). N_female_ ≥ 69 seeds analyzed from each cross with a female T1 parent; N_male_ crosses ≥ 86 seeds from each cross with a male T1 parent. Statistical analyses performed using chi-squared test, comparing the observed numbers of GFP+ and GFP-seeds to 50% GFP+ seeds (1:1 GFP+:GFP-), which is expected if transmission is not affected by the transgene insertion. * represents chi-squared p-value < 0.05.

**Supplemental Figure 4.**
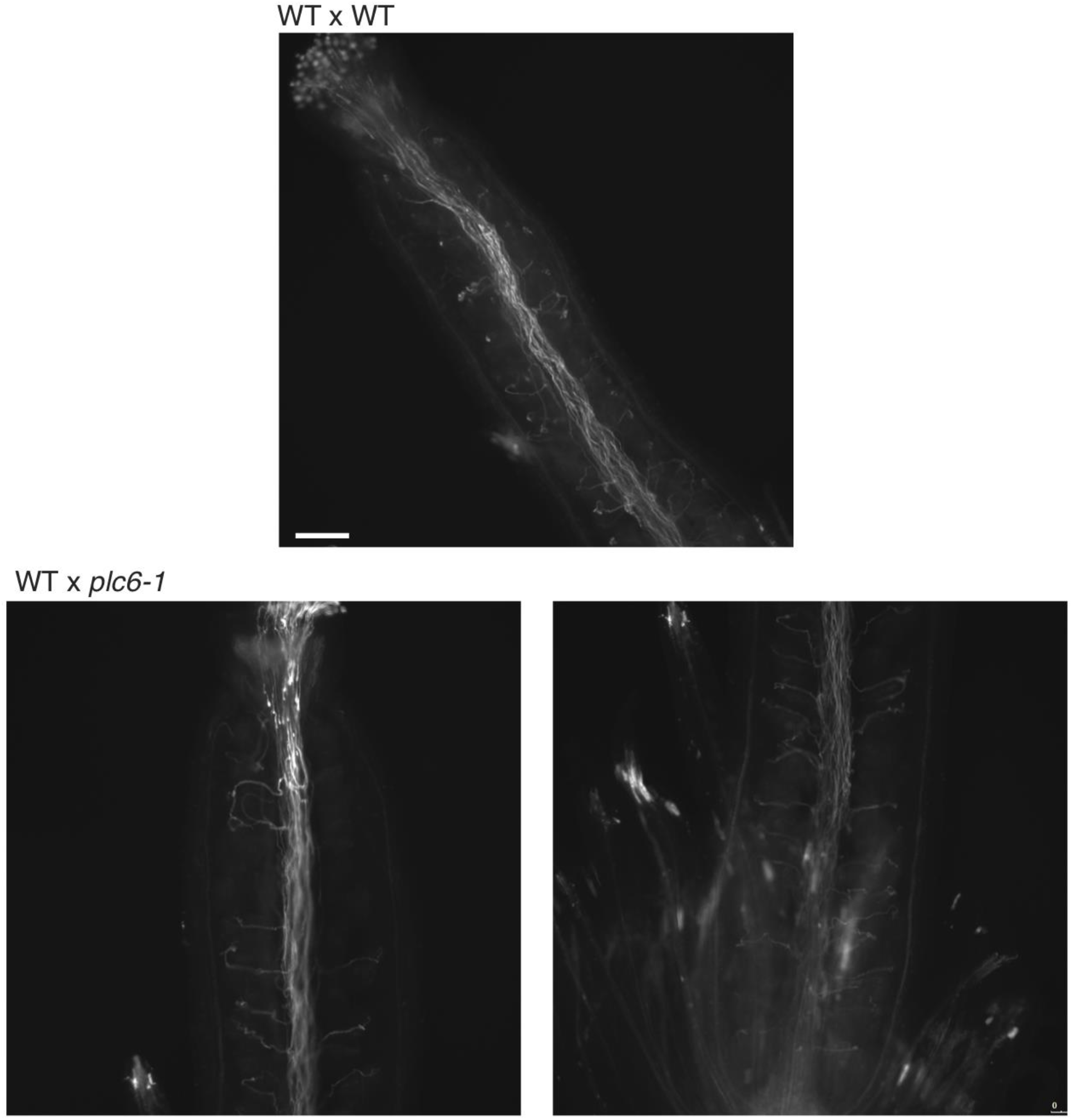
Aniline blue staining of WT and *plc6-4* PTs *in vivo*. WT pistils imaged 24 hours after manual pollination with WT (top) or *plc6-4* (bottom) pollen. Imaged at 10X magnification and scale bar represents 100 μm.

**Supplemental Figure 5.**
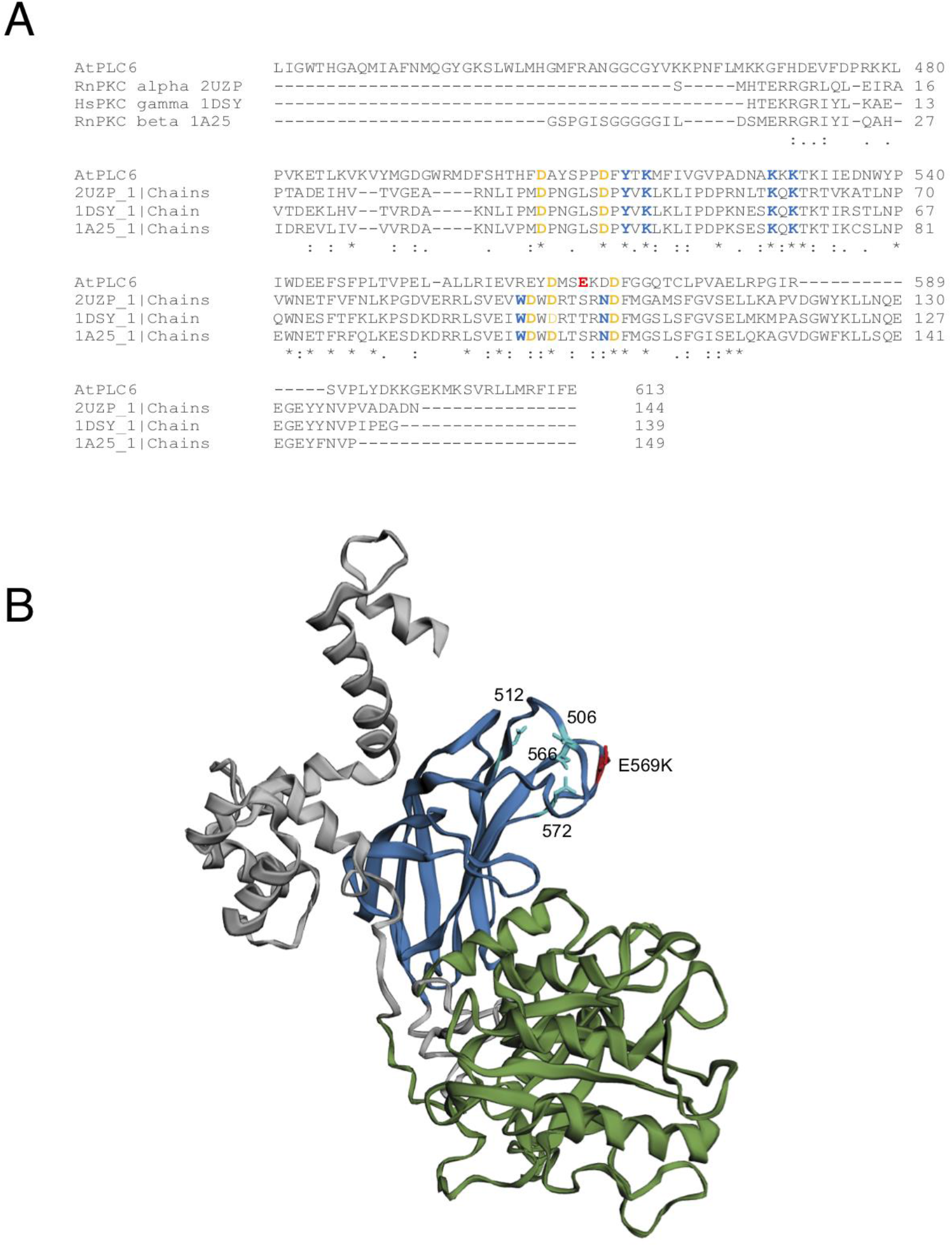
Protein structure of PLC6 protein. Blue is the C2 domain, green is the X and Y catalytic domains, and gray is the EF hand domain plus N-terminal domain of unknown function. A) Alignment of PLC6 C2 domain with the C2 domains of protein kinase Cs from human and rat. Yellow aspartic acid residues are involved in calcium coordination, and blue residues are important for interacting with phosphoinositides. B) 3D protein structure model of PLC6 with calcium-coordinating aspartic acid residues indicated, along with E569.

**Supplemental Figure 6.**
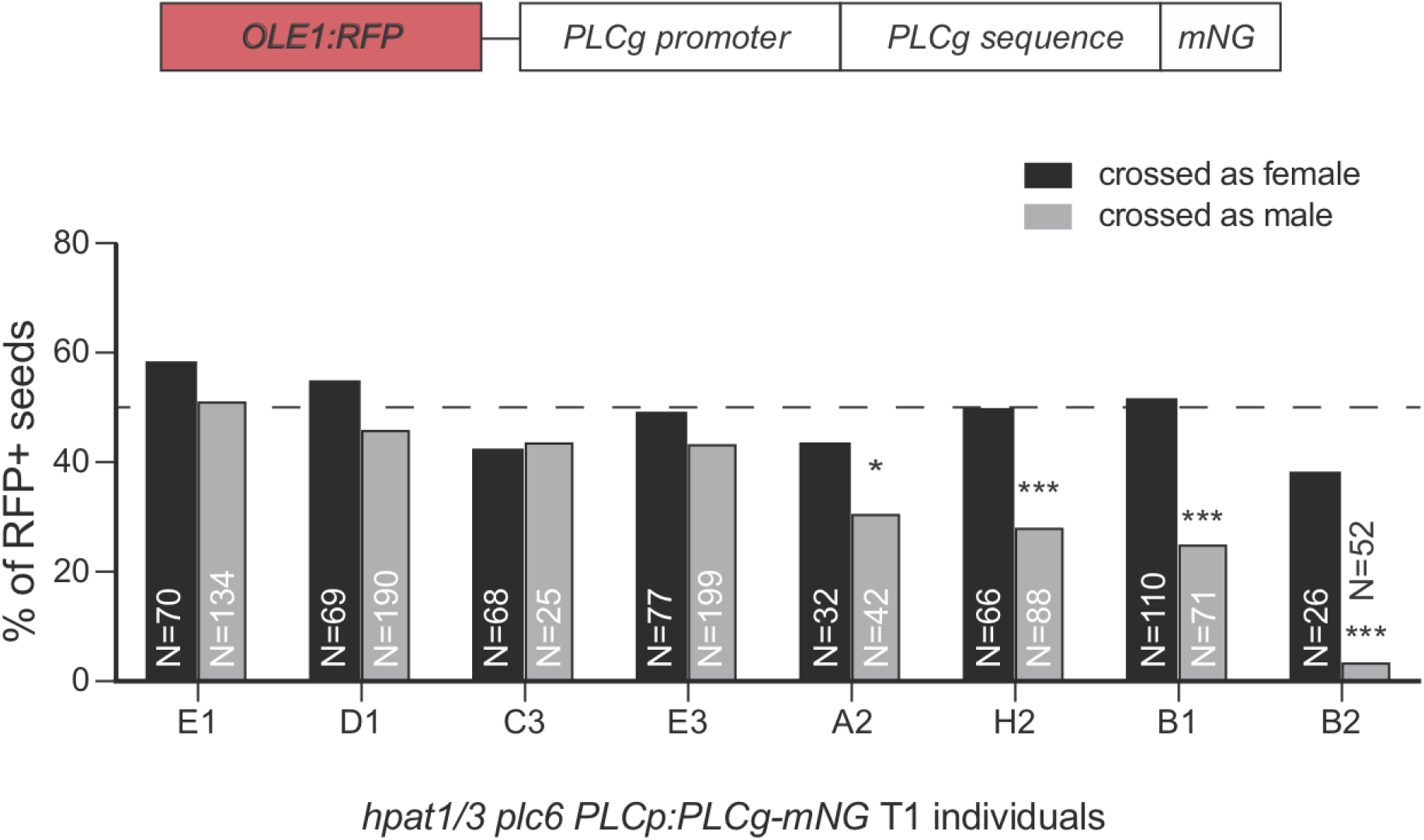
Transmission rates of *PLC6p:PLC-mNG* in *hpat1/3; plc6-4* background. Top) Schematic of PLC6p:PLC6-mNG construct used for expression in WT PTs shown in Figure 8. Bottom) Reciprocal crosses showing %RFP seeds recovered when crossed with WT either as male or female. Statistical analysis performed using chi-squared test.

**Supplementary Table 1.**
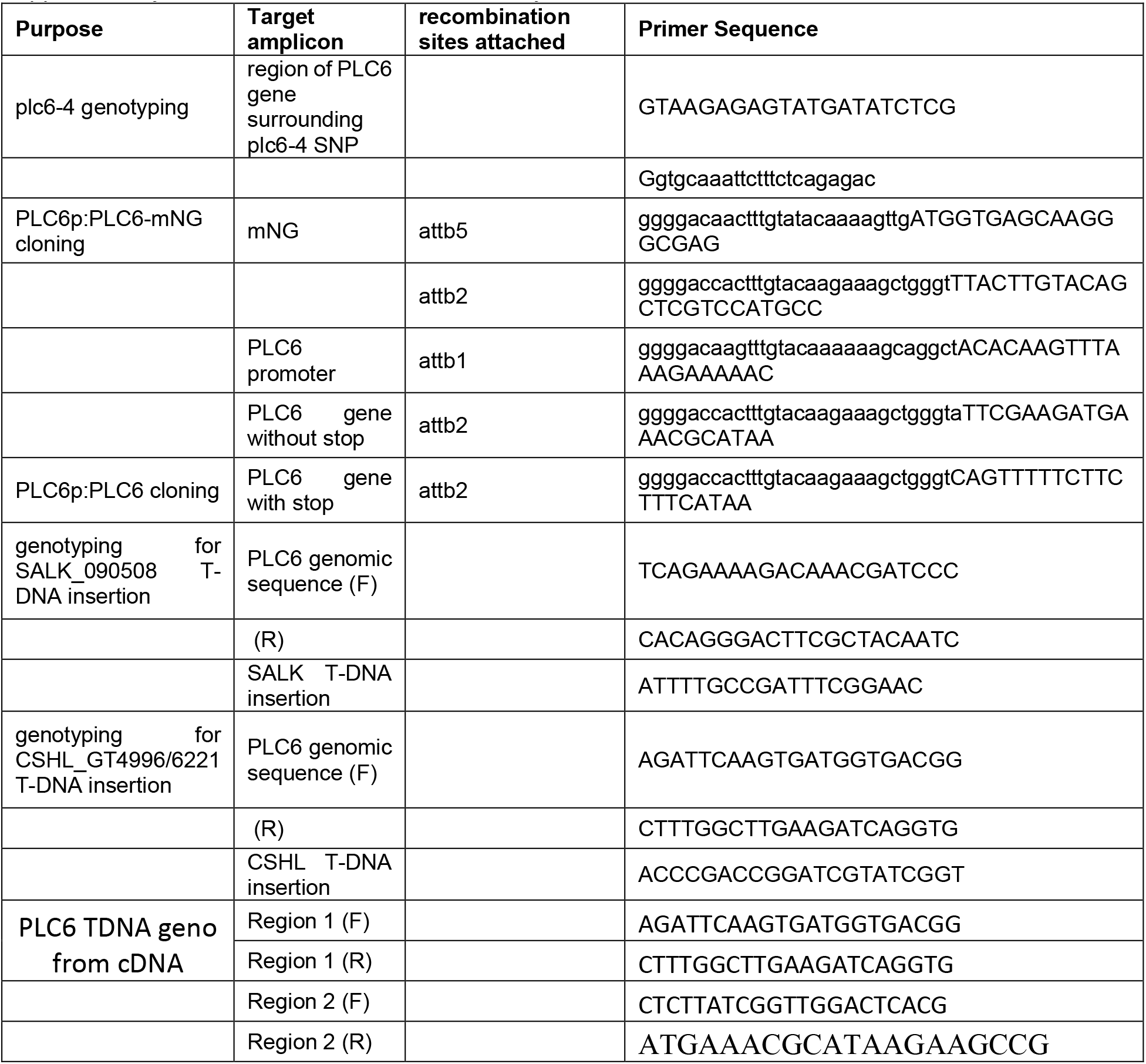
Primers used in this study.

**Supplementary Table 2.**
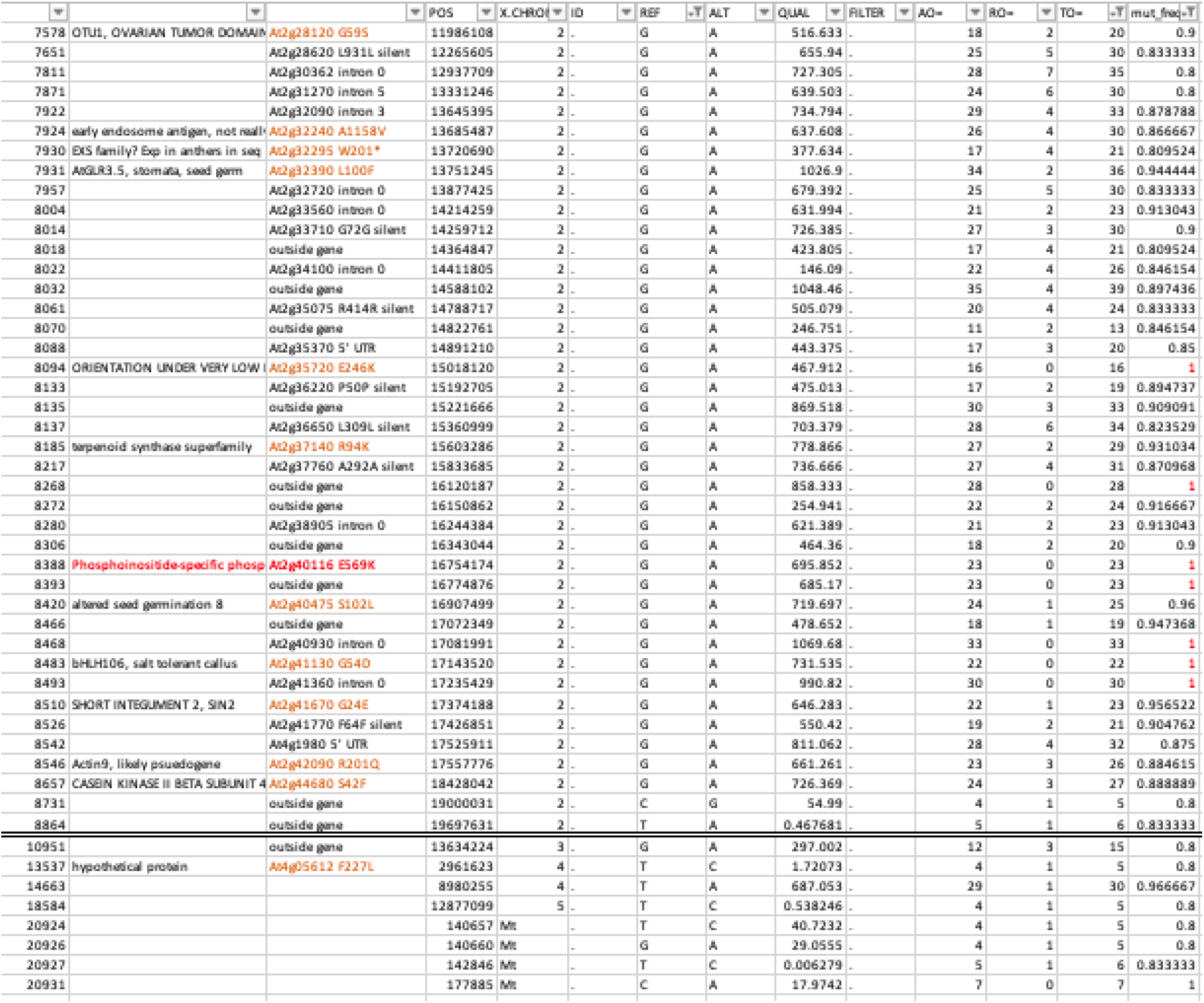
Filtered variants from whole genome sequencing of frh3.

**Supplementary Table 3.** PT lengths after growth on media with different calcium concentrations.

|  | low Ca (1 mM) |  | standard Ca (2 mM) |  | high Ca (4 mM) |  | very high Ca (8 mM) |  |
| --- | --- | --- | --- | --- | --- | --- | --- | --- |
|  | WT | *plc6-4* | WT | *plc6-4* | WT | *plc6-4* | WT | *plc6-4* |
| average (μm) | 236.70 | 162.51 | 252.86 | 136.80 | 257.38 | 119.92 | 165.02 | 70.27 |
| std dev (μm) | 109.20 | 62.48 | 93.94 | 51.42 | 98.41 | 41.20 | 79.94 | 31.72 |
| T-test vs. WT |  | 8.01E-19 |  | 2.21E-49 |  | 3.16E-61 |  | 7.69E-49 |
| T-test vs.<br>std Ca | 0.077 | **6.94E-07** |  |  | 0.60 | **5.67E-05** | **2.28E-26** | **1.62E-51** |

**Supplementary Table 4.**
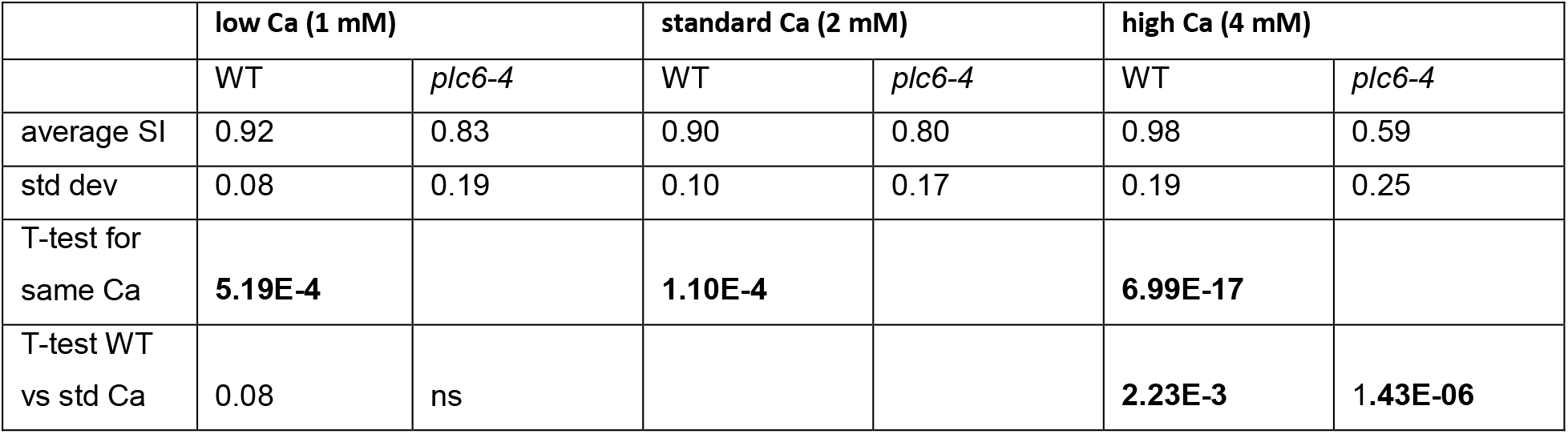
PT straightness after growth on media with different calcium concentrations.

|  | low Ca (1 mM) |  | standard Ca (2 mM) |  | high Ca (4 mM) |  |
| --- | --- | --- | --- | --- | --- | --- |
|  | WT | *plc6-4* | WT | *plc6-4* | WT | *plc6-4* |
| average SI | 0.92 | 0.83 | 0.90 | 0.80 | 0.98 | 0.59 |
| std dev | 0.08 | 0.19 | 0.10 | 0.17 | 0.19 | 0.25 |
| T-test for<br>same Ca | **5.19E-4** |  | **1.10E-4** |  | **6.99E-17** |  |
| T-test WT<br>vs std Ca | 0.08 | ns |  |  | **2.23E-3** | **1.43E-06** |

